# The Tolman-Eichenbaum Machine: Unifying space and relational memory through generalisation in the hippocampal formation

**DOI:** 10.1101/770495

**Authors:** James CR Whittington, Timothy H Muller, Shirley Mark, Guifen Chen, Caswell Barry, Neil Burgess, Timothy EJ Behrens

## Abstract

The hippocampal-entorhinal system is important for spatial and relational memory tasks. We formally link these domains; provide a mechanistic understanding of the hippocampal role in generalisation; and offer unifying principles underlying many entorhinal and hippocampal cell-types. We propose medial entorhinal cells form a basis describing structural knowledge, and hippocampal cells link this basis with sensory representations. Adopting these principles, we introduce the Tolman-Eichenbaum machine (TEM). After learning, TEM entorhinal cells include grid, band, border and object-vector cells. Hippocampal cells include place and landmark cells, remapping between environments. Crucially, TEM also predicts empirically recorded representations in complex non-spatial tasks. TEM predicts hippocampal remapping is not random as previously believed. Rather structural knowledge is preserved across environments. We confirm this in simultaneously recorded place and grid cells.

**One Sentence Summary:** Simple principles of representation and generalisation unify spatial and non-spatial accounts of hippocampus and explain many cell representations.

## Introduction

Humans and other animals make complex inferences from sparse observations and rapidly integrate new knowledge to control their behaviour. Tolman argued that these facilities rely on a systematic organisation of knowledge called a cognitive map (*1*). In the hippocampal formation, during spatial tasks, individual neurons appear precisely tuned to bespoke features of this mapping problem (*2–4*). However, hippocampus is also critical for non-spatial inferences that rely on understanding the relationships or associations between objects and events - termed relational memory (*5*). Whilst it has been suggested that relational memory and spatial reasoning might be related by a common mechanism (*6*), it remains unclear whether such a mechanism exists or how it could account for the diverse array of apparently bespoke spatial cell types.

One promising approach casts spatial and non-spatial problems as a connected graph, with neural responses as efficient representations of this graph (*7, 8*). This has led to new potential interpretations for place cells (*7*) and grid cells (*8, 9*). However, such approaches cannot account for the rapid inferences and generalisations characteristic of hippocampal function in both spatial and relational memory, and do not explain the myriad types of spatial representations observed, or predict how they will change across different environments (remapping).

We aim to account for this broad set of hippocampal properties by re-casting both spatial and relational memory problems as examples of structural abstraction (*10*) and generalisation (Figure 1A-C). Spatial reasoning can be cast as structural generalisation, as different spatial environments share the common regularities of Euclidean space that define which inferences can be made, and which shortcuts might exist. For example, moving *south → east → north → west* will return you to where you started. Structural regularities also permit inferences in non-spatial relational problems. For example, transitive inference problems (which depend on hippocampus (*11, 12*)) require stimuli to be represented on an abstract ordered line, such that *A > B* and *B > C* implies *A > C*. Similarly, abstraction of hierarchical structure permits rapid inferences when encountering new social situations.

**Figure 1:**
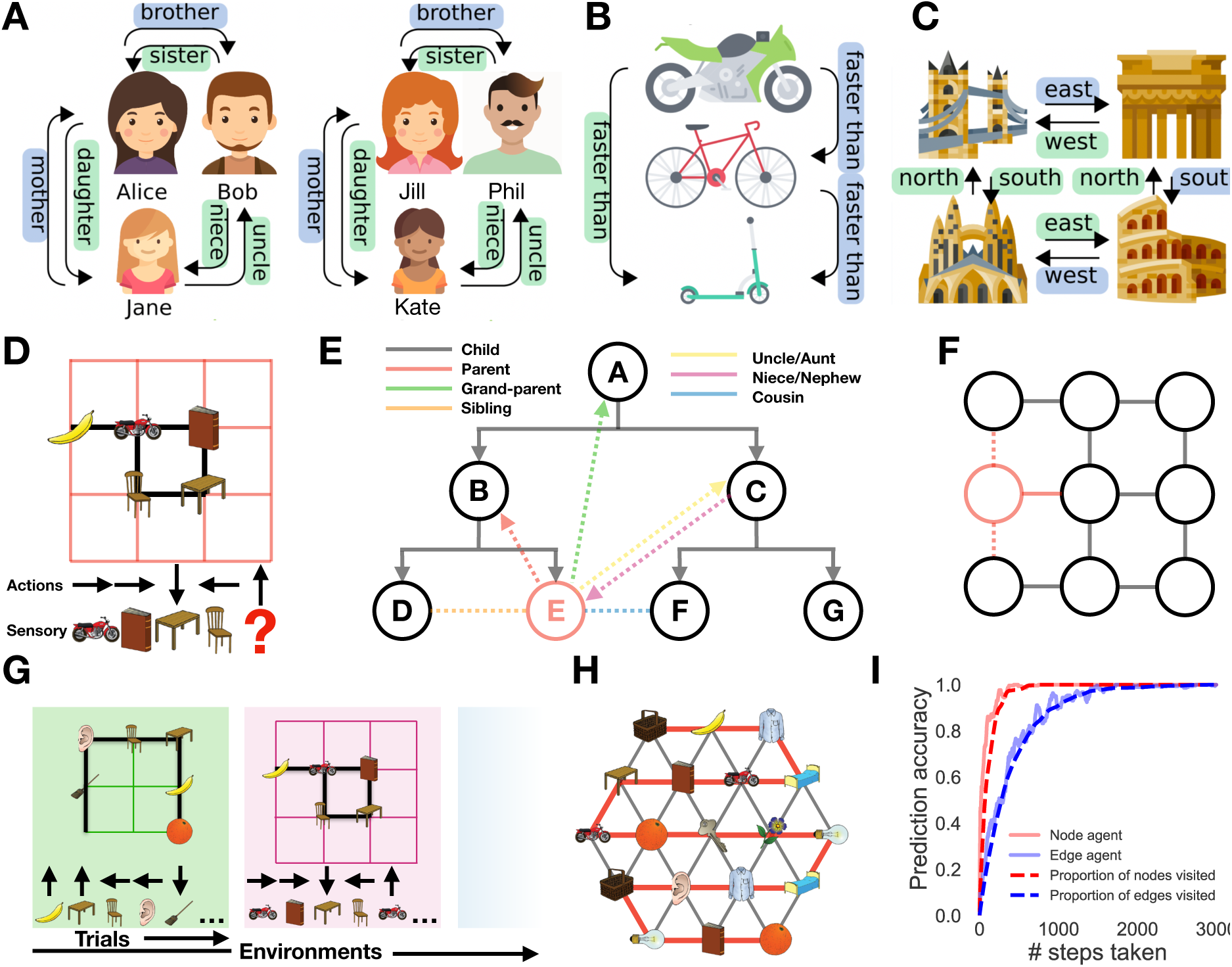
Spatial and relational inferences can be cast as structural generalisation. Structured relationships exist in many situations, and can often be formalised on a connected graph e.g. **A)** Social hierarchies, **B)** transitive inference and **C)** spatial reasoning. Often the same relationships generalised across different sets of sensory objects. This transferable structure allows quick inference, e.g. seeing only the blue relationships allows you to infer the green ones. **D)** Our task is predicting the next sensory observation in sequences derived from probabilistic transitions on the graph. We use arbitrary sensory experiences on each node, e.g. a banana. An agent transitions on the graph observing only the immediate sensory stimuli and associated action taken. For example having seen *motorbike → book → table → chair*, it should predict the *motorbike* next if it understands the rules of the graph. **E)** Should you know the underlying structure of social hierarchies, when observing a new node (in red) via a single relationship - e.g. Emily is Bobs daughter - immediately allows inference about the new nodes (Emilys) relationship to all other nodes (shown in black/gray). **F)** Similarly for spatial graphs observing a new (red) node on the left (solid red line) also tells us whether it is above or below (dashed red lines) other surrounding nodes. **G)** Our agent performs this next step prediction task in many different worlds which share the same underlying structure (e.g. 6- or 4-connected graphs), but they differ in their size and arrangement of sensory stimuli. The agent must learn the commonality amongst all the worlds (the shared graph structure) in order to generalise and perform quick inferences. **H)** Knowing the structure allows full graph understanding after only visiting all nodes and not all edges; here only 18 steps were taken (red line) but a perfect agent infers all 42 links. **I)** An agent that knows structure (node agent) will reach peak predictive performance after it has visited all nodes; quicker than one that has to see all transitions (edge agent). Icons made by Nikita Gobulev (transport), FreePik (faces) and Abib Sulthon (landmarks) from flaticons.com.

Structural generalisation offers dramatic benefits for new learning and flexible inference, and is a key issue in artificial intelligence. One promising approach is to maintain factorised representations in which different aspects of knowledge are represented separately and can then be flexibly re-combined to represent novel experiences (*13*). Factorising the relationships between experiences from the content of each experience could offer a powerful mechanism for generalising this structural knowledge to new situations. Notably, exactly such a factorisation exists between sensory and spatial representations in medial and lateral entorhinal cortices respectively. Manns and Eichenbaum propose that novel conjunctions of these two representations form the hippocampal representation for relational memory (*14*).

We demonstrate that this factorisation and conjunction approach is sufficient to build a relational memory system (the Tolman-Eichenbaum machine; TEM) that generalises structural knowledge in space and non-space; predicts a broad range of neuronal representations observed in spatial and relational memory tasks; and accounts for observed remapping phenomena in both hippocampus and entorhinal cortex. Notably, although hippocampal remapping is thought to be random, TEM predicts that this apparent randomness hides a structural representation that is preserved across environments. We verify this prediction in simultaneously recorded place and grid cells. These results suggest a general framework for hippocampal-entorhinal representation, inference and generalisation across spatial and non-spatial tasks.

## Results

### Spatial and relational inferences can be cast as structural generalisation

We consider the unsupervised learning problem where an agent must predict the next sensory experience in a sequence derived from probabilistic transitions on graphs (Figure 1D). The agent does not see the graph, only a sequence of sensory observations and the action or relation that caused each transition. Different types of relation exist, e.g. in a family hierarchy, parent, aunt, child and nephew imply different transitions on the graph, but each transition-type has the same meaning at every point on the graph. Similarly, in space, action is defined by heading direction (e.g. NESW on 4-connected graphs).

If all transitions have been experienced, the graph can be stored in memory and perfect predictions made without any structural abstraction. However, if structural properties of the graph are known a priori, perfect prediction is possible long before all transitions have been experienced; it only requires each node to have been experienced (Figure 1H,I): This can be easily understood: When the structure of the graph is known, a new node can be introduced with a single relation (Bob has a daughter, Emily - Figure 1E) and all other relations can immediately be inferred (Emily is Alices granddaughter and Cats niece etc.). Similarly, in space, if the structure of 2D graphs is known, then placing a new node on an X-Y coordinate is sufficient to infer relational information to every other point on the graph (Figure 1F).

In summary, after experiencing many graphs with different sensory observations and learning their common relational structure, an agent should maximise its ability to predict the next sensory observation after each transition on a new graph (Figure 1G).

### The Tolman Eichenbaum machine

To build a machine that solves this problem, we first consider a normative solution. We then show that a tractable approximation to this solution maps simply onto the functional anatomy of the hippocampal system. We give complete descriptions of TEM in the SOM, at both intuitive and detailed levels of description.

### Generative model

We want to estimate the probability of the next sensory observation given all previous observations on this and all other graphs. A parsimonious solution will reflect the fact that each task is composed of two factors, a graph-structure and sensory observations (Figure 2A).

**Figure 2:**
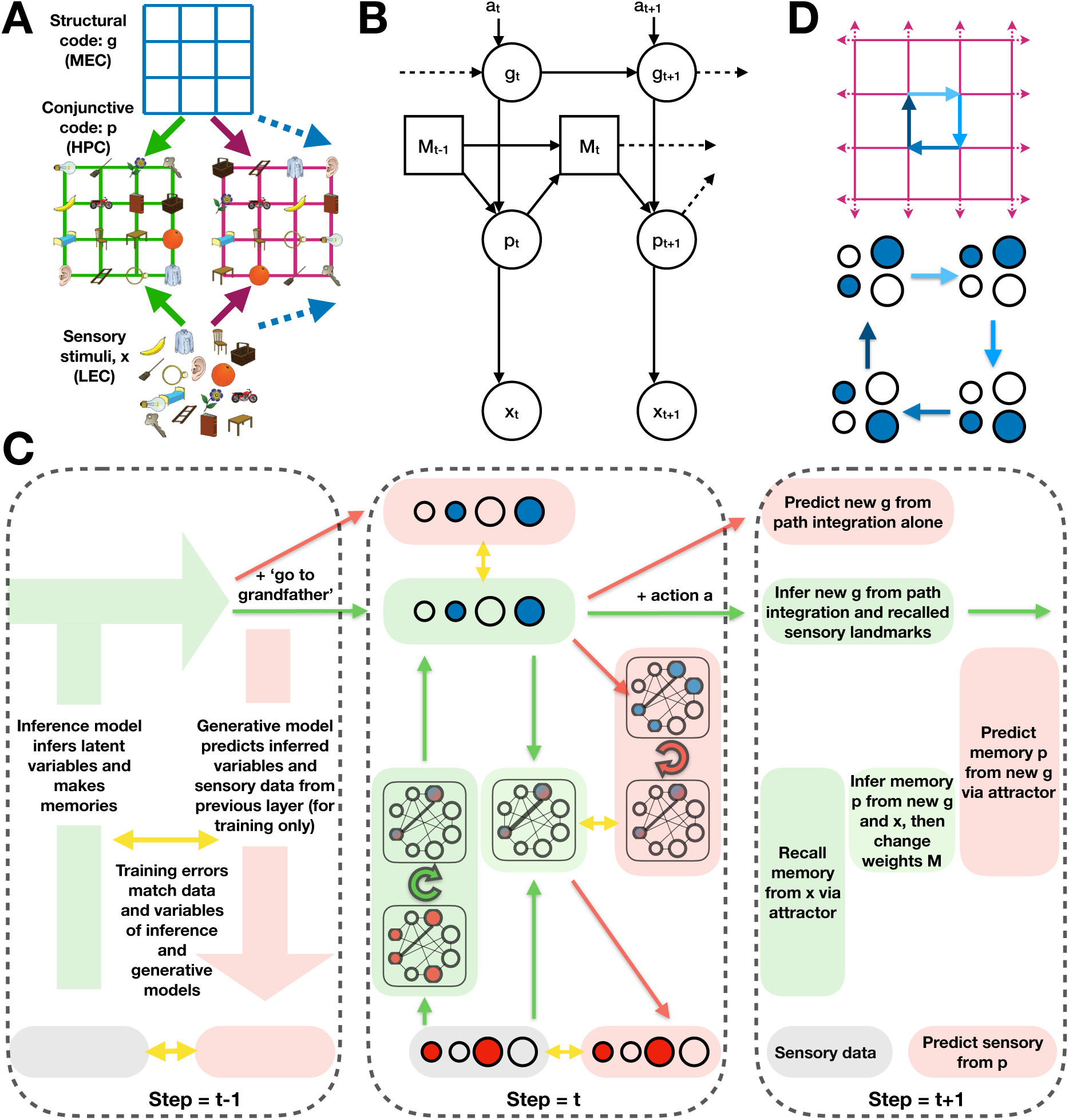
The Tolman-Eichenbaum Machine. **A)** Factorisation and conjunction as a principle for generalisation. Separating structural codes (the transition rules of the graph) from the sensory codes (the sensory objects on each node) allows generalisation over environments that share the same structure. The conjunctive code represents the current environment in the context of this learned structure. **B)** The generative model of TEM with observed variables **x***_t_* and **a***_t_* (sensory experience and action taken respectively), and latent variables **g***_t_* and **p***_t_* (where I am and a memory of what is where respectively). This is a graphical model which shows the dependencies of variables, not the connections. See Figure 2C (red) for equivalent connections. TEM transitions through latent variables **g**, and retrieves memories **p** using Hebbian weights M. **C)** Depiction of TEM at three time-points, with each time-point described at a different level of detail. Green/ red show inference and generative networks. Timepoint *t* − 1 shows the overall Bayesian logic, t shows network implementation, *t* + 1 describes each computation in words. Circles depict neurons (blue is **g**, red is **x**, blue/red is **p**); shaded boxes depict computation steps; arrows show learnable weights (green and red are weights in inference and generative networks); looped arrows describe recurrent attractor. Black lines between neurons in attractor describe Hebbian weights M. Yellow arrows show training errors. **D)** TEM must learn structural codes that 1) have a different representation for each state so that different memories can be stored and retrieved, but also 2) have the same code on returning to a state (from any direction) so the appropriate memory can be retrieved.

This problem can be expressed graphically as a generative model (Figure 2B), where latent variables are positions that result from taking relational actions in a cognitive map. To facilitate generalisation of knowledge across domains we separate latent variables of abstract location that generalise across maps, **g**, from those that are *grounded* in sensory experience and therefore specific to a particular map **p**. Each variable is represented as a population (vector) of units. Conditioned on an action, **g** transitions to a new abstract location. This new **g** can then index Hebbian memories M, retrieving a memory of grounded location **p**, that, if the state has been visited before, can correctly predict the sensory experience **x**.

This *generative model* allows sensory data to be simulated from an arbitrary map but, to be useful, we must also be able to *infer* the latent variables [**p**, **g**] from real sensory data, so that predictions can be made about **this** map. In Bayesian reasoning, this involves ***inverting*** the generative model. The exact solution to this problem is intractable, so we turn to approximate methods.

### Inference and the hippocampal formation

We use modern Bayesian methods (*15, 16*) to *learn* an inference network (Figure 2C, green) that maps the stream of observations and actions to the latent variables, updating abstract location, **g**, and grounded location, **p**, as new sensory data appear. This inference network is independent of the generative model, but, once trained, approximates optimal inference on the generative models latent variables (See **Model training** and SOM).

Although the parameters of the inference network are learned, we are able to make critical choices in its architecture. The resulting network maps simply onto the functional anatomy of the hippocampal formation and its computations and can be intuitively represented in schematics (Figure 2C, green).

Following Eichenbaum (*14*), hippocampal representations, (**p**), are inferred as a conjunction between sensory input (**x**) in the lateral (LEC), and abstract location (**g**) in the medial (MEC) entorhinal cortex. Mirroring hippocampal synaptic potentiation (*17*), this enables memories to be rapidly stored in weights (M) between **p** using simple Hebbian learning between co-active neurons; and retrieved by the natural attractor dynamics of the resultant auto-associative network (Figure 2C middle).

To infer a new **g** representation, TEM performs path integration from the previous g, conditional on the current action/relation. This can be related to continuous attractor models (CANNS, (*18*)) of grid cells (*19*). Like CANNs, different recurrent weights mediate the effects of different actions/relations in a recurrent neural network (Figure 2C middle). Unlike CANNs, however, weights are not hardcoded, but **learnt from experience**, allowing map-like abstractions and path integration to extend to arbitrary non-spatial problems.

Path integration accumulates errors (*20*). To overcome this problem TEM can take advantage of a second source of information about **g** - the conjunctive representations, **p**, stored in the hippocampal memory M. TEM indexes M with the current sensory experience, **x**, to retrieve a set of candidate representations of **g** (previously visited places with a similar sensory experience), and uses these to refine the path integrated **g** via a Bayesian update.

When representing tasks that have self-repeating structure, it is efficient to organise cognitive maps hierarchically. To allow such hierarchy to emerge, we separate our model into multiple parallel streams, each as described above. These streams are only combined when retrieving **p** in the attractor network (see SOM for details).

### Model training

Both the generative and inference models have weights that must be learnt. The objective of training is for the generative model to *predict the sensory input*, x, and for the inference model to *infer the generative models latent variables*, [**p**, **g**], from the sensory input. The resulting training algorithm (Figure 2C) involves an interplay between generative (red) and inference (green) models, in which the generative model takes the current state of the inference model and (from this) predicts its next state (including the next sensory data). This leads to errors between the predicted and inferred/observed variables at each level [**p**, **g**, **x**]. The weights in both networks are adjusted along a gradient that reduces these errors using backpropagation through time (see SOM).

This scheme is similar to the Wake Sleep algorithm (*21*) and Helmholtz Machine, which learn generative and inference models for sensory inference (*22*). Unlike these sensory models through, TEM predicts temporal sequences (*16*) and uses an attractor memory to allow rapid generalisation of structural knowledge. The reliance on generative model predictions is notable as hippocampal replay appears to sample from a generative model of the environment (*23, 24*).

The model is trained in multiple different environments, differing in size and sensory experience. Different environments use the same network weights, but different Hebbian weights, M. The most important weights are those that transition **g** as they encode the structure of the map. They must ensure 1) that each location in the map has a different **g** representation (so a unique memory can be built), 2) that arriving at the same location after different actions causes the same **g** representation (so the same memory can be retrieved) - a form of path integration for arbitrary graph structures. For example, the relation uncle must cause the same change in **g** as father followed by brother, but different from brother followed by father.

### TEM generalises structural knowledge to novel sensory environments

We first test TEM on classic non-spatial relational memory tasks thought to depend on hippocampus - transitive inference and social hierarchy tasks. After training, TEM makes first presentation inferences in novel transitive inference tasks (Figure 3F) e.g. regardless of particular sensory identities (e.g. *A, B, C, D, E* or *cat, dog, elephant, fox, badger*) after being shown sequences such as *A > B > C > D > E*, TEM answers B to the query what is 3 more than E, i.e. TEM has learned ordinal structural knowledge. Equally, in social hierarchy tasks, TEM infers relationships based on limited information (Figure 3E) e.g. after being shown that Bob is the brother of Cat who is Frans mother, TEM answers Fran when queried who is Bobs niece?. In both cases, although TEM had never seen the particular sensory details of the task before, it had seen similar relational structures which it could learn from and generalise. Such first presentation inferences we refer to as zero-shot inference - *only* possible with learned structural knowledge.

**Figure 3:**
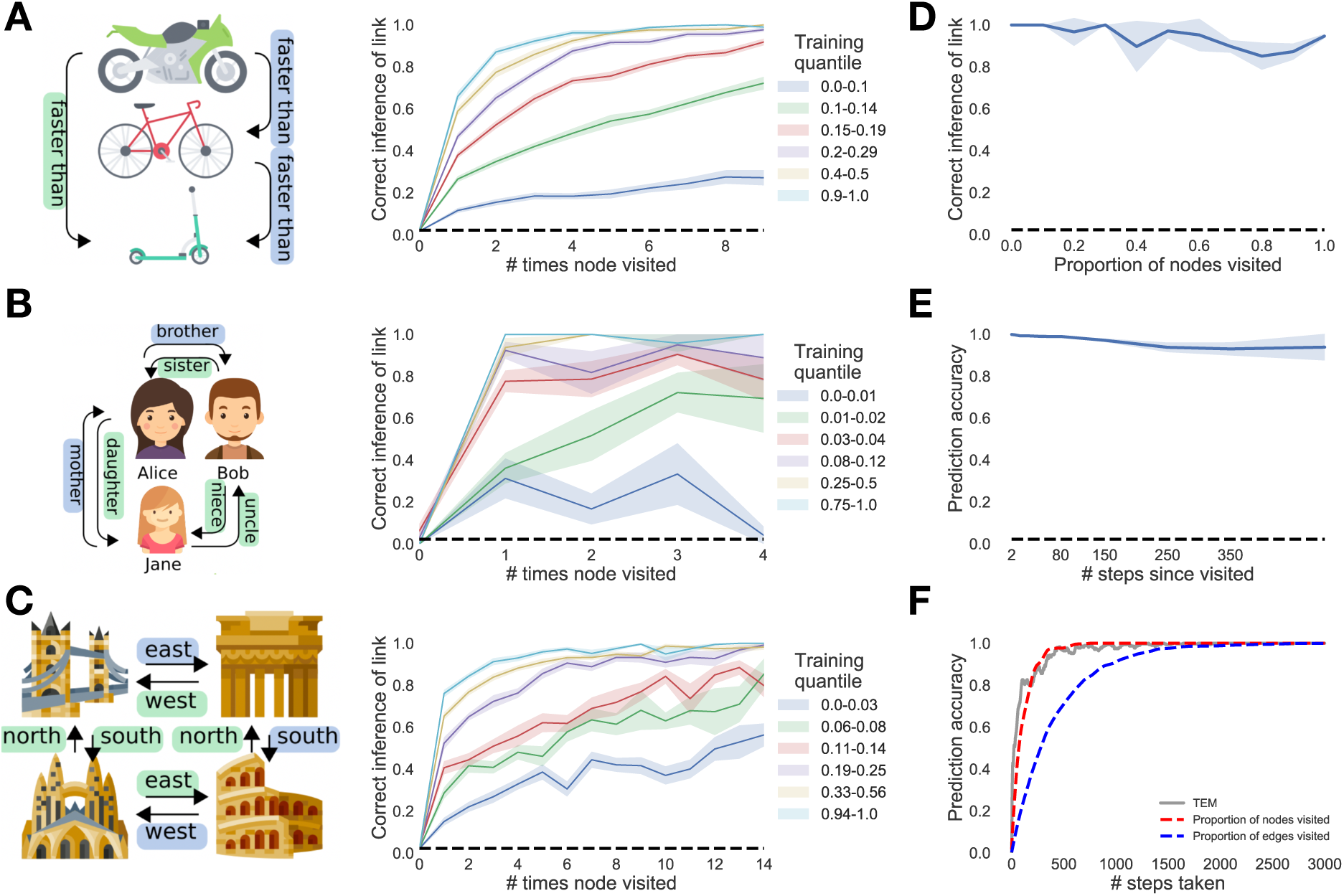
TEM learns and generalises abstract relational knowledge. **A-C)** Learning to learn: When TEM has only seen a few environments (blue/green) it takes many visits to each node for it to be remembered - this is because it 1) does not understand the structure the graph and 2) it has not learned to how utilise memories. After visiting more environments and discovering the common structure (cyan/yellow) TEM is able to correctly predict a node on the second visit regardless of the edge taken - TEM now understands both the rules of the graph and how to store and retrieve memories. **A)** Transitive inference graphs, **B)** social hierarchy graphs and **C)** 2D spatial graphs. **D-F)** On 2D graphs. **D)** Here we consider how TEM performs on zero-shot link inference: TEM is asked to predict sensory observations when returning to a node via a *new* direction - this is only possible with learned structural knowledge. TEM is able to do this. **E)** TEM is able to store memories for long periods of time, even though it is only trained on sequences of length 25. **F)** TEMs performance tracks nodes visited not edges. All these results demonstrate that TEM has learned and generalised abstract structural knowledge.

Knowing the underlying structure allows one to know the entire relational structure after a single visit to each state (Figure 1H,I). TEM demonstrates this data efficiency with its performance in line with the proportion of states visited in the graph, not the edges taken (Figure 3D).

We also test TEM on tasks with an underlying spatial structure (e.g. Figure 1F,H). Again, TEM performs zero-shot inference in spatial generalisation tasks (Figure 3C) - only possible with learned structural knowledge, whilst also storing long term relational memories (Figure 3B).

### TEM represents structure with grid cells that generalise across environments

We begin by considering TEM agents that diffuse from state to state at random on 2D graphs, constrained only by the neighbourhood transitions in the environment. Here, TEMs abstract location (**g**) representations resemble grid cells (Figure 4A) and band cells (Figure 4B) as recorded in rodent medial entorhinal cortex (*4, 25–27*). As in the brain, we observe modules of grid cells at different spatial frequencies and, within module, we observe cells at different grid phases (Figure 4A).

**Figure 4:**
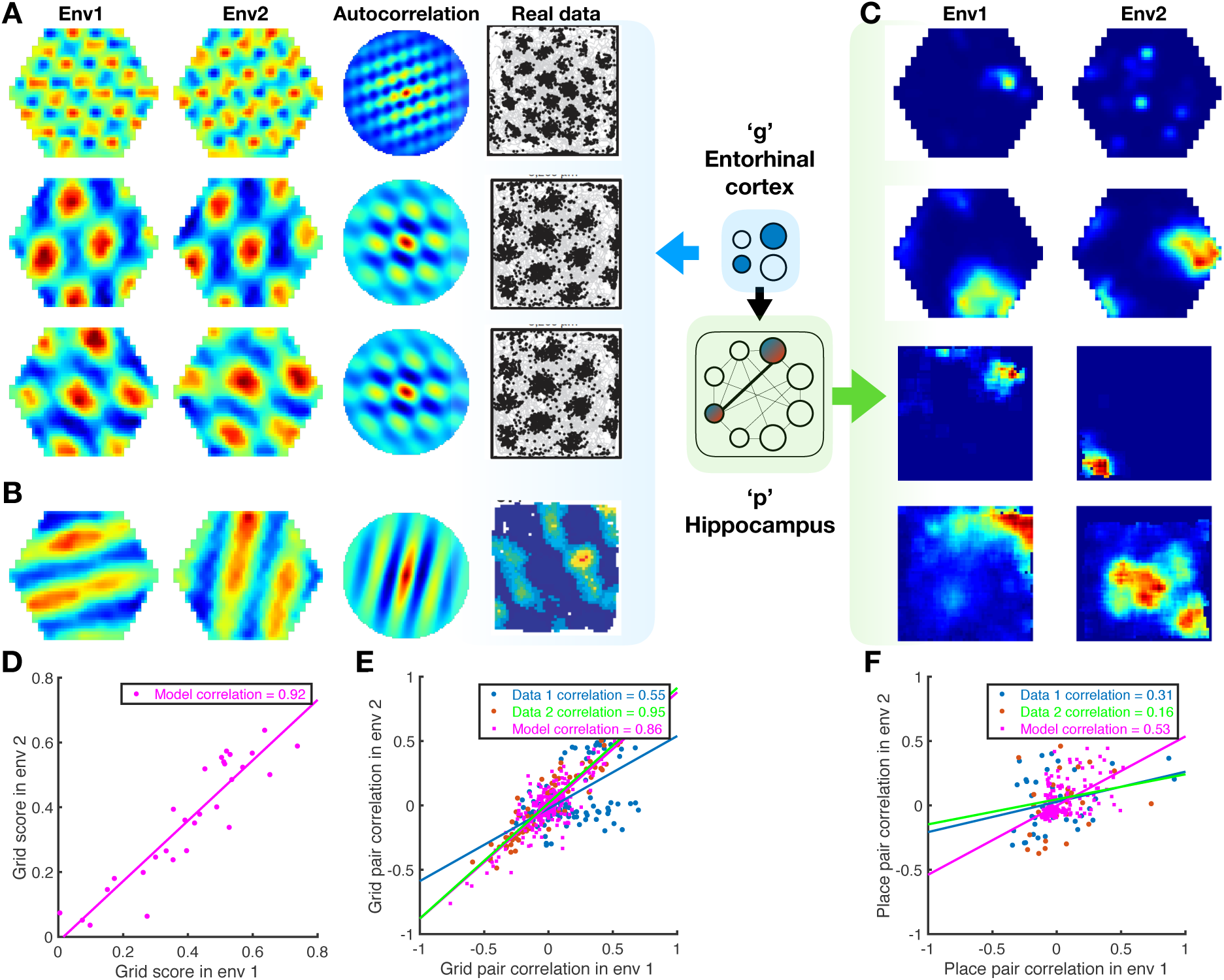
TEM learns structural grid cells that generalise and conjunctive memory place cells that remap. **A-B)** TEM learned structural representations for random walks on 2D graphs (left to right: environments 1 and 2, autocorrelation, real data from (*25, 34*), different cells shown vertically). **A)** TEM learns grid-like cells, of different frequencies (top vs middle), and of different phases (middle vs bottom). **(B)** Band like cells are also learned by TEM. Importantly these TEM structural representations A-B) generalise across the different environments 1 and 2. **(C)** Learned memory representations resemble place cells (left/right: environments 1 and 2; top 2 simulated, bottom 2 real cells) and have different field sizes. These cells remap between environments (left/right), i.e. do not generalise. **D)** Grid scores of TEM grid-like cells correlate across environments. **E-F)** To examine whether relationships between cells are preserved between environments, we correlated the spatial correlation coefficients of pairs of grid or place fields from each environment, using data from (*35*), and (*36*). **E)** The spatial correlation coefficients of pairs of TEM structural cells and real data grid cells correlate strongly. **F)** TEM hippocampal and real data place cells preserved their relationship to a lesser extent. This suggests that TEM structural cells, along with real grid cells, encode generalisable relationships to a greater extent that TEM hippocampal and real place cells.

TEMs top-level (**g**) representations reflect the need to be both maximally different at different spatial locations, to enable independent memories at each location and invariant to approaching the same location from different trajectories (path integration) so that the correct memory can be retrieved. Our results suggest that these two constraints are sufficient to produce grid- and band- like representations.

Importantly, top-layer TEM representations generalise, retaining their properties across different environments. This is true of both the first and second-order properties of the population. For example, a grid cell in environment 1 is a grid cell of the same frequency in environment 2, and the correlation structure across grid cells is preserved - that is grid cells (in the same module) that fire next to each other in one environment, do so in all environments. This generalisation is agnostic to environment size, thus TEM has not just learned to represent a single environment size but has instead learned a general representation of 2D topology. These preserved properties provide the substrate for generalisation of relational structure, and are also observed in rodent grid cell populations recorded in multiple environments (*28, 29*).

### TEM forms memories with place cell representations that remap across environments

In TEM, each hippocampal cell, **p**, is a conjunction between a TEM structural entorhinal cells, **g**, and sensory input, **x**; they will only be active when both the structural cells and sensory input are both active (Figure S4). In diffusive worlds, TEM learns sparse representations that resemble hippocampal place cells (Figure 4C). These place-like fields span multiple sizes, mirroring the hierarchical composition of hippocampal place fields (*30, 31*).

Importantly TEMs hippocampal cells, unlike their entorhinal counterparts, do not generalise. Although each environment shares the same structure, the sensory objects have a different distribution. The conjunctive nature of the hippocampal representation means that TEMs hippocampal cells *do not fully* preserve their correlation structure across environments (Figure 4F) but instead relocate apparently at random in different environments. This phenomenon is commonly observed in rodent hippocampal cells and is termed *global remapping* (*32, 33*).

### Diverse Entorhinal and hippocampal cell types form a basis for transition statistics

Animals do not move by diffusion (*37*). We next examined representations learned by TEM when trained with different behavioural paradigms (Figure 5, see SOM for full experimental details). For non-diffusive transitions, we mimic animals that prefer to spend time near boundaries, and approach objects. Here, because the transition statistics change, so do the optimal representations for predicting future location. Indeed, training TEM on these *behavioural* transition statistics leads to the emergence of new cellular representations that are also found in rodents. Now entorhinal representations, **g**, in TEM include border cells (*38*) (Figure 5C) and cells that fire at the same distance and angle from any object (object vector cells (*39*); Figure 5A) for the two cases respectively. This is easily understood: In order to make next-state predictions TEM learns predictive representations, with object vector cells predicting the next transition is towards the object - as is behaviourally the case.

**Figure 5:**
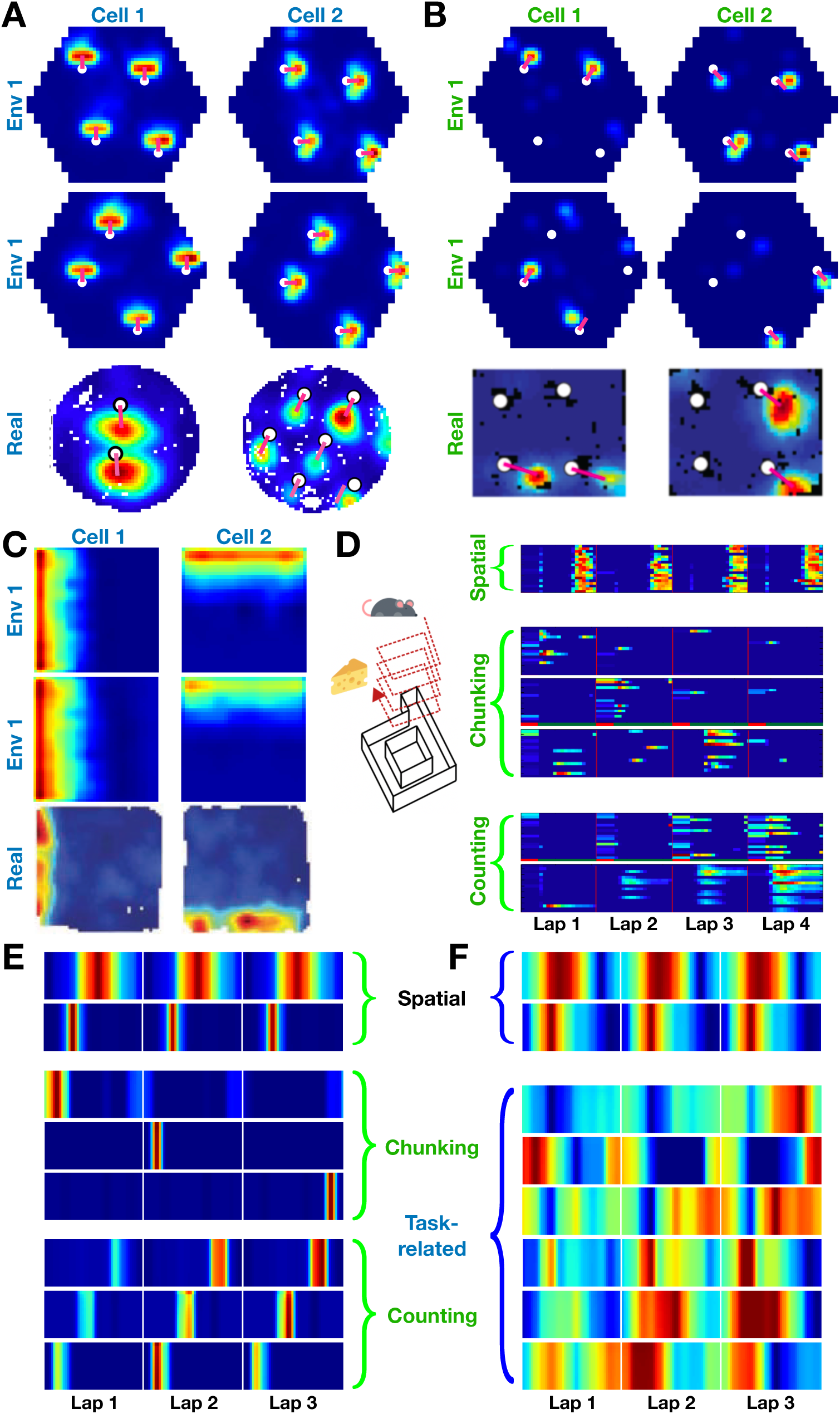
TEM learned representations reflect transition statistics. When the agents transition statistics mimic different behaviours TEM learns different representations (left to right: different cells, top to bottom: environments 1 and 2 and real data). **A)** When biased to move towards objects (white dots) TEM learns structural cells that are active at a certain distance and orientation from the object - object vector cells (*39*). These cells generalise to *all* objects. **B)** TEM hippocampal cells also reflect this behavioural transition change with similar cells. However, these TEM hippocampal cells do not generalise to all objects - landmark cells (*40*). **C)** When biased to stay near boundaries, TEM is more likely to learn border cell like representations. **D)** In (*41*), rodent performs laps of a track, only rewarded every 4 laps. Different hippocampal cell types are found: spatial place like cells (top), those that preferentially fire on a given lap (middle) and those that count laps (bottom). **E)** When TEM performs a similar task, only rewarded every 3 laps (see SOM), similar representation are learned. **F)** TEM entorhinal cells not only learn spatially periodic cells (top), but also cells that represent the complex non-spatial task structure of every 3 laps (bottom). Cells of this type are yet to have experimentally observed, but are predicted by TEM. Icons made by FreePik (mouse) and Smashicons (cheese) from flaticons.com.

Critically, these TEM entorhinal cells also generalise across environments, with TEM object vector cells generalising to all objects both within and across environments. The cells do not represent the objects themselves, but rather their predictions about transitions, and they do so in a way that generalises, allowing immediate inferences in new environments. Notably, these same properties are observed in object vector cells in rodent entorhinal cortex (*39*) (Figure 5A).

Similar cells exist in TEMs hippocampal layer, **p**, with a crucial difference. Here, object sensitive cells represent the vector to a particular object-type but do not generalise across objects (Figure 5B) - they represent the conjunction between the structural representation and the sensory data. These cells are reminiscent of landmark cells that have been recorded in rodent hippocampus (but not entorhinal cortex) (*40*).

Objects occur at random locations within each environment, thus when representing the transition statistics of the environment, TEM’s entorhinal layer **g** is able to arbitrarily *compose* object vector cell representations (at any location) along with grid and other entorhinal representations. These results suggest that the zoo of different cell types found in entorhinal cortex may be viewed under a unified framework - summarising the common statistics of tasks into basis functions that can be flexibly *composed* depending on the particular structural constraints of the environment the animal/agent faces.

### A mechanistic understanding of complex non-spatial abstractions

While cellular responses are well understood in rodent open field tasks, we have little knowledge of how they combine to control behaviour in more complex problems. Because TEM can learn arbitrary structural abstractions, it can also account formally for hippocampal and entorhinal responses in complex non-spatial tasks.

To illustrate this, we consider a recent finding by Sun et al. (*41*). Rodents perform laps of a circular track, but only receive reward every four laps. Now hippocampal cells develop a new representation. Whilst some cells represent location on the track, (i.e. place cells; Figure 5D top), others are also spatially selective, but fire only on one of the 4 laps (Figure 5D middle). A third set fire again at a set spatial location, but vary their firing continuously as a function of lap number (Figure 5D bottom). Hippocampal cells maintain a complex combined representation of space and lap number.

When TEM was trained on this task (using 3 laps instead of 4 for computational reasons), it learned these same 3 representations in hippocampus (Figure 5E, see SOM for further cells). Here “reward” was just a particular sensory event that repeated every 3 trials. As in the biological data, some TEM hippocampal cells encode location on every lap. These cells allow TEM to predict the sensory events that are unchanged between laps. However, as in recorded data, some TEM cells encode location on of the three laps, and some with a lap gradient. These cells allow TEM to represent its position within the 3-lap cycle.

Importantly, TEM allows us to reveal a candidate mechanism. TEMs medial entorhinal cells have reconfigured to code differently for each lap (Figure 5F top), understanding that the abstract task space is not a single lap but three (Figure 5F bottom). This mechanism is consistent with empirical data as manipulation of entorhinal cortex removes lap-sensitive hippocampal cells (*41*). However, TEMs entorhinal representations stand as a prediction as Sun et al. did not record entorhinal cells.

These results suggest entorhinal cells can learn to represent tasks at multiple levels of cognitive abstraction simultaneously, with hippocampal cells reflecting their conjunction with sensory experience. More broadly, TEM predicts hippocampal cells will reflect a combination of the necessity to predict the current sensory input and the necessity to separate states that predict different distant futures. They will therefore contain representations of the current location in space, but also the current location in the task.

### Structural knowledge is preserved over apparently random hippocampal remapping

The critical assumption that enables TEMs structural inferences is that the hippocampal representations of new events are **not random**. Instead they are constrained by learned structural representations in the entorhinal input. This assumption seems at odds with the commonly held belief that hippocampal place cells remap randomly between environments. However, the structural representation for space is *periodic*. Thus, place cells can preserve structural information across environments without being spatial neighbours with the same cells in each environment. Instead, individual cells need only to retain their phases with respect to the grid code. Here, structural knowledge is retained but remapping still occurs because place cells might, in a new environment, move to the same phase but with respect to a **different grid peak** (see e.g. Figure 6A). Together with the different sensory input between environments, this leads to remapping in TEMs conjunctive hippocampal cells (Figure 4C).

**Figure 6:**
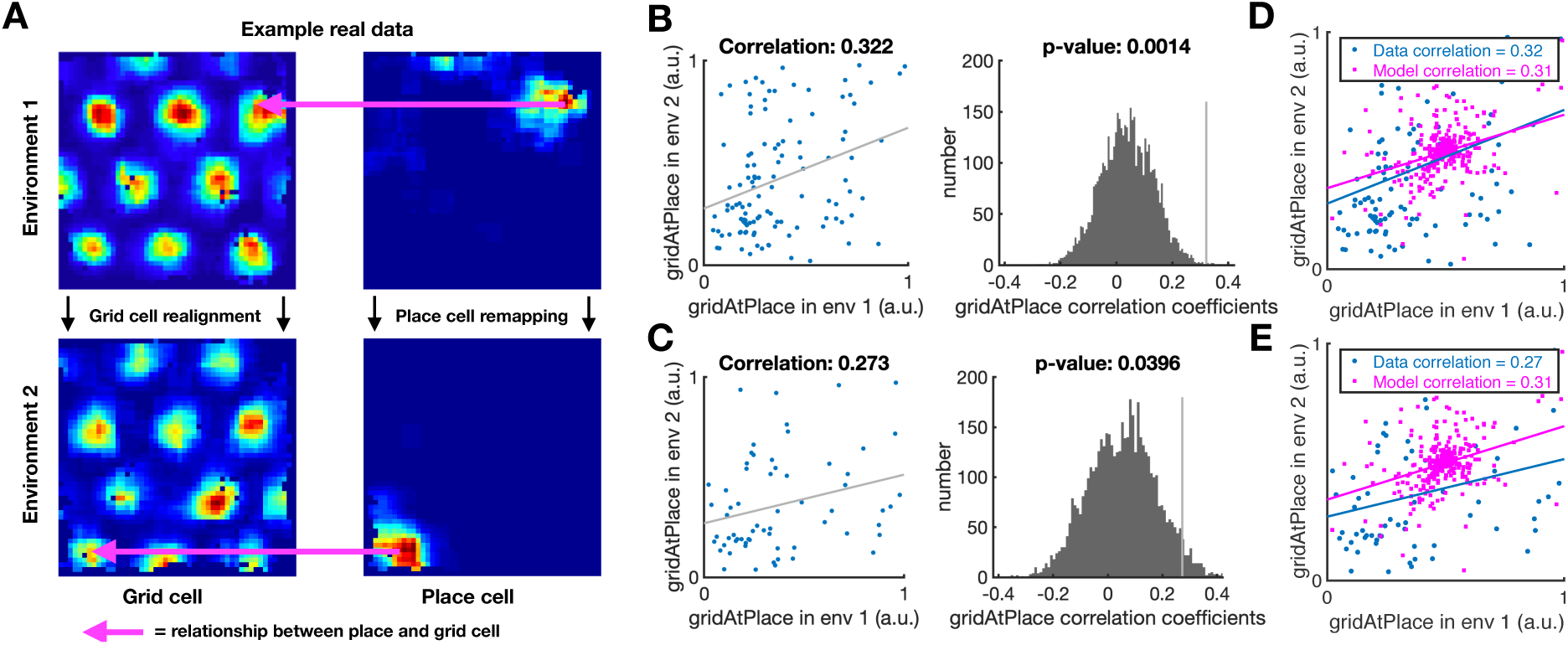
Structural knowledge is preserved over apparently random hippocampal remapping. TEM predicts that place cells remap to locations consistent with a grid code. **A)** A place cell co-active with a certain grid cell will be more likely to remap to locations that are also co-active with that grid cell. **B-C)** Open field remapping experiments with simultaneously recorded place and grid cells (*35, 36*). We compute the grid cell firing rate at the location of place cell peak for every grid cell - place cell pair in each of the two environments, and then correlate this measure across environments (left) (see SOM for details). We compare this correlation coefficient to those computed from the same procedure with randomly permuted place cell peaks (right). This is done for two independent datasets B) Barry et al. (*35*) and **C)** Chen et al (*36*). The true observed correlation coefficients like off the null distribution (p¡0.05), demonstrating that place cell remapping is not random, but instead tied to structural constraints of grid cells. **D-E)** The same analysis on TEM learned representations shows qualitatively similar results.

Is this true in biological remapping? We tested data from two experiments in which both place and grid cells have been recorded whilst rats (*35*) and mice (*36*) freely forage in multiple environments. Experiment 1 (*35*) has two environments of the same dimensions (1m by 1m) but differing in their sensory cues so the animals could distinguish between them. Each of seven rats has recordings from both environments. Twenty-minute recordings are taken each day in both environments. Experiment 2 (*36*) has four mice in a real and a virtual reality environment of sizes (60cm by 60cm). Trials in the real and virtual environments were 20 and 40 minutes long respectively.

We asked if the activity of each grid cell at the peak firing of each place cell peak (gridAtPlace) is correlated across environments. If the place cell fires at the same grid phase in each environment then the same grid cell will have strong activity at each place cell peak. To test the significance of this correlation, we perform a permutation test by generating a null distribution from randomly shifted place cells.

For experiment 1 we find significant correlation for the 115 within-animal place cell-grid cell pairs that satisfy conservative criteria (see SOM) (*r* = 0.322, *p <* 0.01 (permutation test) - Figure 6B) and for the liberal set of 255 pairs (*r* = 0.63, *p <* 0.05 - SOM). We replicate these results in dataset 2 across 64 conservative pairs (*r* = 0.273, *p <* 0.05 - Figure 6C) and the liberal set of 148 pairs (*r* = 0.544, *p <* 0.05 - SOM). These results are robust to all combinations of parameters settings for our criteria of cell acceptance to the analysis (see SOM). We also show that a second measure is significant for experiment 1 and trending for experiment 2 (SOM).

If there were only a single grid frequency (or module) in entorhinal cortex, we would expect a near perfect correlation across environments between gridAtPlace scores for each grid-cell place-cell pair. Whilst both datasets have non-zero correlations, the correlation is far from perfect (Figure 6). This would be expected if either (1) place cells are influenced by phases of more than a single grid module or (2) place cells predominantly received input from a single grid module, but we (the experimenter) do not know which module. Therefore, in order to gauge the magnitude of the effect, we performed the same analysis on TEM representations. Data and model show similar correlations (average *r_data_* = 0.27 *−* 0.32, *r_model_* = 0.31) (Figure 6D,E).

To our knowledge, this is the first data demonstration of non-random place cell remapping, and support a key prediction of our model: that hippocampal place cells, despite remapping across environments, retain their relationship with the entorhinal grid, providing a substrate for structural inference.

## Conclusion

Building an understanding that spans from computation through cellular activity to behaviour is a central goal of neuroscience. One field that promises such an understanding is spatial navigation and the hippocampus. However, whilst cells are precisely described for open field foraging in small arenas, it has been unclear how these responses could generalise to real-world behaviours. Similarly, it has been unclear how to understand these spatial responses in the context of hippocampal involvement in memory broadly writ (*42*) and relational memory in particular (*6*). In this work, we have shown that by formalising the problem of relational abstraction, using factorisation and conjunction of representations, it is possible to account for spatial inferences as a special case of relational memory as hypothesized by Eichenbaum and colleagues

In doing so, we provided unifying principles that account for a number of seemingly disparate phenomena. For example, grid cells, band cells, border cells, object vector cells all emerge from bases describing likely transitions. We show that this structural basis is also important for understanding several seemingly unrelated processes such as hippocampal remapping (*32, 33*), and transitive inference (*11*) which are shown to be two sides of the same coin - the structural knowledge transferred during remapping supports the generalisation of transitive structure. Whilst the idea that hippocampal memories are summarised in cortex is influential (*43*), TEM therefore also suggests how cortical representations feed back onto the hippocampus to organise new experience and memory.

Spatial reasoning not only provides a particularly clean example of the factorisation of relational and sensory knowledge, but also affords generalisation as relational meaning repeats regularly across space (*2*). However, by considering the relational memory problem more broadly, we have shown that cellular responses can be formally accounted for in situations more complex than open-field foraging. Whilst we have so far considered simple behaviours (e.g. running 4 laps of a maze to attain reward), we envisage that, together with exciting insights from reinforcement learning (*8*), this and similar frameworks may be useful in extending our computational understanding from open field navigation towards Tolmans original ideas of a systematic organisation of knowledge spanning all domains of behaviour (*1*).

RIP Howard Eichenbaum (1947-2017).

## Acknowledgements

We would like to thank Jacob Bakermans for help with figure preparation, and to Philip Schwartenbeck and Sebastian Vasquez-Lopez for providing helpful comments.

## Funding

We would like to thank the following funding sources. EPSRC scholarship to JCRW, MRC scholarship to THM, Wellcome Senior Research Fellowship (104765/Z/14/Z) and JS McDonnell Foundation award (JSMF220020372) to TEJB, Wellcome Collaborator award (214314/Z/18/Z) to TEJB, NB, CB. Wellcome Principle Research Fellowship (202805/Z/16/Z) and ERC Advanced Grant NEUROMEM to NB. Wellcome Senior Research Fellowship (212281/z/18/z) to CB. The Wellcome Centre for Integrative Neuroimaging and Wellcome Centre for Human Neuroimaging are each supported by core funding from the Wellcome Trust (203139/Z/16/Z, 203147/Z/16/Z).

## Author contributions

JCRW developed the model, performed simulations and drafted paper. CB, GC and NB collected data. JCRW, THM analysed data. JCRW, TEJB conceived project with input from THM, SM initially. JCRW, TEJB edited paper with input from all other authors.

## Competing interests

Authors declare no competing interests.

## Data and materials availability

All code will be released on publication.

## Supplementary Materials

### 1 Task details

We wish to formalise a task type that not only relates to known hippocampal function, but also tests the learning and generalising of abstract structural knowledge. This would then offer a single common framework (generalising structural knowledge) that explains many hippocampal functions, and as it turns out, explains many known cell representations.

We formalise this via relational understanding on graph structures (a graph is a set of nodes that relate to each other). We demonstrate that two seemingly distinct hippocampal functions - spatial navigation and relational memory tasks - can be viewed in this common framework.

Should one passively move on a graph (e.g. Figure S1A), where each node is associated with a non-unique sensory observation (e.g. an image of a banana), then predicting the subsequent sensory observation tests whether you understand the graph structure you are in. For example, if you return to a previously visited node (Figure S1A pink) by a new direction - it is only possible to predict correctly if you know that a *right → down → left → up* means you’re back in the same place. Knowledge of such loop closures is equivalent to understanding the structure of the graph.

We thus train our model (TEM) on these graphs with it trying to predict the next sensory observation. TEM is trained on many environments sharing the same structure, e.g. 2D graphs (Figure S1A), however the stimulus distribution is different (each vertex is randomly assigned a stimulus). Should it be able to learn and generalise this structural knowledge, then it should be able to enter new environments (structurally similar but with different stimulus distributions) and perform feats of loop closure on first presentation.

We note that this feat of loop closure on first presentation has an intuitive meaning for space, but it is identical to the first presentation inferences made on tasks of transitive inference (*11*) and social hierarchy (*44*) - tasks that the hippocampus plays a crucial role in.

In order to show that under a single framework (TEM), many aspects of hippocampus and entorhinal cortex can be explained, we thus choose graphs structures that reflect the types of tasks (space, transitive inference, social hierarchies) under which these brain areas have been studied. We describe the details of each task, along with details of simulations in Section 4.

### 2 The Tolman-Eichenbaum Machine

In the following description, we try to repeat the same information at successively increasing levels of detail. We hope this will allow readers to build an understanding of the model at their preferred level.

#### 2.1 Problem statement - Intuitive

We are faced with the problem of predicting sensory observations that come from probabilistic transitions on a graph. The training data is a continuous stream of sensory observations and actions/relations (Figure S1A). For example, the network will see banana, north, tree, east, book, south, door or Joe, parent, Denise, niece, Anna, sibling, Fred. The model should predict the next sensory observation with high probability.

#### 2.2 Problem statement - Formal

Given data of the form 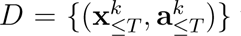 with *k* ∈ {1, …, *N*} (which environment it is in), where **x***_≤T_* and **a***_≤T_* are a sequence of sensory observations and associated actions/relations (Figure S1A), *N* is the number of environments in the dataset, and *T* is the duration of time-steps in each environment, our model should maximise its probability of observing the sensory observations for each environment, p*_θ_* (**x***_≤T_*), where *θ* are the model parameters.

#### 2.3 High level model description

We choose to model our problem probabilistically using a generative model - this allows us to offer a normative model for how the observed data depends on unobserved latent variables e.g. seeing a banana depends on where you are, but where you are is a latent variable - it is never observed. One can then in principle use Bayes theorem to invert this generative model and provide the optimal posterior distribution over the latent variables given the observations (inference). However, in most scenarios, including ours, this inversion is computationally impractical. Many methods have been proposed to **approximate** this inference. One particularly powerful method is to learn the parameters of an inference model. Once trained, this model will approximately invert the generative model and perform the inference, mapping the observed data to the latent (unobserved) variables. This idea was introduced in the Wake-sleep algorithm (*21*) and the Helmholtz machine (*22*), and has since been adopted by Variational Autoencoders (*15, 45*).

In common instantiations of generative models, latent variables are the causes of, for example, pixels in stationary images. Here, we provide a generative model where latent variables are positions that result from taking relational actions in a cognitive map. We further enable generalisation of knowledge across domains by separating latent variables of location that generalise across maps, **g**, from those that are grounded in sensory experience and therefore specific to a particular map **p**. Grounded locations, **p** encode [abstract location, sensory experience] conjunctions for the current environment.

The model aims to predict the next sensory experience from all previous sensory experiences. This problem is not inherently *Markovian*. The next sensory experience can depend on historic experiences independently from the most recent experience (old locations might be encountered on meandering paths). However, the problem can be rendered Markovian by the inclusion of a Memory M that remembers what experience is where in the current environment. The inclusion of grounded location variables, **p**, means that this memory simply has to remember these variables.

We give the full generative model for the general probabilistic case with noise in both the action and sensory inputs and derive the appropriate loss functions. However, in this manuscript we only consider tasks where there is no noise in these inputs. We therefore implement a version of the model that ignores noise (discussed in Section 3) - this leads to faster training and more accurate inference in the noise-free case.

#### 2.4 TEM and the brain

We propose TEM’s abstract location representations (**g**) as medial entorhinal cells, TEM’s grounded locations (**p**) as hippocampal cells, and TEM’s sensory input **x** as lateral entorhinal cells. In other words, TEM’s sensory data (the experience of a state) comes from the ‘what stream’ via lateral entorhinal cortex, and TEM’s abstract location representations are the ‘where stream’ coming from medial entorhinal cortex. TEM’s (hippocampal) conjunctive memory links ‘what’ to ‘where’, such that when we revisit ‘where’ we remember ‘what’.

TEMs medial entorhinal representation, (**g**), invites comparison to continuous attractor neural networks (CANNs) (*18*), commonly used to model grid cell activity in spatial tasks (*19, 46, 47*). Like CANNs, TEMs **g** layer is a recurrent neural network. Like CANNs, different recurrent weights mediate the effects of different actions/relations. Unlike CANNs, however, our weights are not hardcoded, but learnt from experience. Furthermore, due to the factorisation afforded by **p**, they can be learnt directly from sensory experience without any location input. They can therefore learn abstract map-like representations not only in spatial problems, but also in arbitrary non-spatial problems - even those in which it would be difficult for humans to hand code an effective location representation (such as a family tree).

TEM’s grounded locations (**p**) resemble hippocampal cells, encoding location-sensory conjunctions (*48–50*) and enabling fast episodic memories (*32, 51, 52*).

TEMs sensory representations (**x**) resemble lateral entorhinal representations, encoding processed sensory input (here - objects) (*14, 53, 54*). Notably, TEM learns most effectively if sensory representations are passed through an approximate Laplace Transform as is reported in lateral entorhinal cells (*55*) (see section 2.8 and Section 2.10).

TEM describes the hippocampal-entorhinal system as one that performs inference; TEM medial entorhinal cells infer a location in abstract space based based on their previous belief of location (and optionally sensory information linked to previous locations via memory). TEM hippocampal cells infer the current memory representation based on a conjunction between the sensory data and believed location in abstract space.

Though we already refer to these variables as entorhinal and hippocampal cells, we reiterate that no representations are hard-coded - *all TEM representations are learned*.

#### 2.5 High-level algorithmic description

We now describe the fundamentals behind the Tolman-Eichenbaum Machine (TEM). TEM sees a stream of sensory observations and actions (**x** and **a**). Its objective is to predict the next sensory input. If these observations are arranged on a graph with any regularities, TEM can profit from these regularities to predict the sensory consequences of edges it has never previously taken. After learning these regularities, TEM can transfer them to new environments that have the same regularities, but different sensory observations.

##### 2.5.1 Principles

TEM relies on two simple principles / components. Firstly a map-like component that learns about the abstract structure shared across environments (Tolman), and secondly a conjunctive memory component that grounds this learned abstract structure to the current environment (Eichenbaum). We denote the map-like variables as **g**, and the grounded conjunctive variables as **p**.

Each grounded location **p** is a conjunction, tying and abstract location **g** to a sensory experience **x**. Each abstract location, however, has the potential to instantiate many different grounded locations - one for each for possible sensory experience. An attractor network memory learns, after a single experience, which location-experience pair is valid in the current world. The opposite is also true - a sensory experience can re-instantiate the memory of a grounded location i.e. the conjunctive memory process allows both abstract location to predict sensory experience, and sensory experience to predict abstract location.

Naturally, TEM can only predict a sensory observation should it have seen it before and formed a memory of its grounded location. TEM re-instantiates memories of grounded locations via indexing from its abstract location, **g**, and so re-instantiating the correct grounded location requires TEM to index using the same abstract location code as when the memory of grounded location was formed.

This puts strong constraints on the types of representations TEM must learn. Firstly, it must learn a structural map-like code that transferably path-integrates such that **g** is the same when returning to a state (so the correct memory is indexed). Secondly it must learn representations **g** that are different for different states - so that each state can have a separate memory attached to it. These two constraints are fundamental to TEM representations, and are shown to be satisfied by grid-cell and other entorhinal codes.

##### 2.5.2 Generative model

The generative model sees an action **a**, combines this with its previous **g** to predict the next abstract location in its cognitive map **g** which then proposes candidate grounded locations. An attractor network pattern completes these candidates, suppressing those that have not been experienced before, to restore a memory of the appropriate grounded location **p**. The restored memory/grounded location then predicts sensory observation **x**. We show this schematically in Figure S4B

##### 2.5.3 Inference Model

The inference model sees a sensory observation **x**, retrieves a memory of grounded location best related to this sensory observation, then infers the next **g** from both the previous **g** (and action **a**) and this memory of grounded location. **p** is then re-inferred using the best estimate of **g** and the new **x** (Figure S4A). This new grounded location **p** is used to update memory weights, *M*.

##### 2.5.4 Training

Both the generative and inference models have weights that must be learnt. The objective of training is for the generative model to predict the sensory input, **x**, and for the inference model to infer the generative models latent variables, [**p**, **g**], from the sensory input. The resulting training algorithm (Figure 2C) involves an interplay between generative and inference models, in which the generative model takes the current state of the inference model and (from this) predicts its next state (including the next sensory data). This leads to errors between the predicted and inferred/observed variables at each level [**p**, **g**, **x**]. The weights in both networks are adjusted along a gradient that reduces these errors using backpropagation through time.

The model is trained in multiple different environments, differing in size and sensory experience. When entering a new environment, network weights are retained but Hebbian weights M are reset. The most important weights are those that transition **g** as they encode the structure of the map. They must ensure 1) that each location in the map has a different **g** representation (so a unique memory can be built), 2) that arriving at the same location after different actions causes the same **g** representation (so the same memory can be retrieved) - a form of path integration for arbitrary graph structures. For example, the relation uncle must cause the same change in **g** as father followed by brother, but different from brother followed by father (’non-associative’ relationships, not just associative ones seen in 2d graphs). These transition weights are shared between generative and inference models, though other weights are not. Shared weights are atypical for variational autoencoders, but are important for biological considerations. At each step we compared what was inferred to what was predicted from the inferred at the previous time-step - we add up the losses for a sequence and then update the weights.

##### 2.5.5 Hierarchies in the map

When representing tasks that have self repeating structure (as ours do), it becomes efficient to hierarchically organise your cognitive map. In this spirit, we separate our model into multiple parallel streams, each as described above, where each stream can learn weights to represent the world at different scales. These scales are then only combined when retrieving memories/ grounded locations in the attractor network. We provide further details on this is Section 2.10.

##### 2.5.6 Model flow summary

The inference model is the one that sees sensory data **x***_t_* at each time-step *t*. It is awake and transitions through time on its own inferring **g***_t_* and **p***_t_* at each time-step. The inference model infers the new abstract location **g***_t_* before inferring the new grounded location **p***_t_*. In other words latent variables **g** and **p** are inferred in the following order **g***_t_*, **p***_t_*, **g***_t_*_+1_, **p***_t_*_+1_, **g***_t_*_+2_ ⋯. This flow of information is shown in a schematic in Figure S2B.

Independently, at each time-step, the generative model asks are the inferred variables from the inference model what I would have predicted given my current understanding of the world (weights). I.e. 1) Is the inferred **g***_t_* the one I would have predicted from **g***_t−_*_1_. 2) Is the inferred **p***_t_* the one I would have predicted from **g***_t_*. 3) Is **x***_t_* what I would have predicted from **p***_t_*. This leads to errors (at each timestep) between inferred and generative variables **g***_t_* and **p***_t_*, and between sensory data **x***_t_* and its prediction from the generative model.

At then end of a sequence, these errors are accumulated, with both inference and generative models updating their parameters along the gradient that matches each others variables and also matches the data.

Since the inference model runs along uninterrupted, its activity at one time-step influence those at later time-steps. Thus when learning (using back-propagation through time - BPTT), gradient information flows backwards in time. This is important as, should a bad memory be formed at one-time step, it will have consequences for later predictions - thus BPTT allows us to learn how to form memories and latent representations such that they will be useful many steps into the future.

#### 2.6 Detailed algorithmic description

#### 2.7 Generative architecture

TEM has a generative model (Figure S2) which factorises as

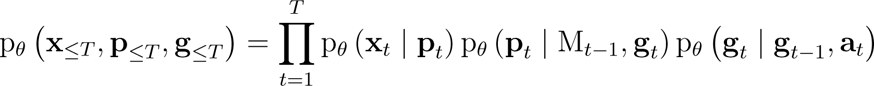

M*_t−_*_1_ represents the agent’s memory composed from past hippocampal representations **p***_<t_*. *θ* are parameters of the generative model. The initial p*_θ_* (**g**_1_ *|* **g**_0_, **a**_1_) = p*_θ_* (**g**_1_), i.e not conditioned on any prior variables. The model can be described by a sequence of computations represented by the following equations:

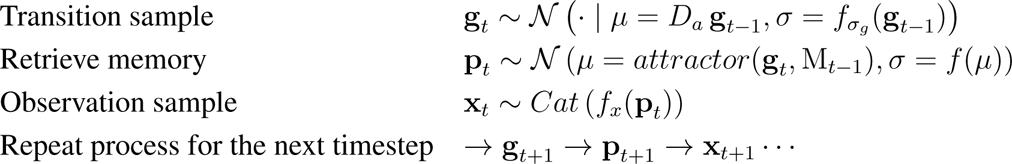

We pictorially show these processes in Figure S5 (just consider the blue freq stream initially, the second red stream will make sense in Section 2.10).

To predict where we will be, we can transition from our current location based on our heading direction (i.e. path integration). *D_a_* is a set of learnable weights for each available action.

Once TEM has transitioned, it then retrieves a memory indexed by it’s believed location. Memories are retrieved via an attractor network (details in a later section). *f_σg_* is a simple multi layer perceptron (MLP).

After the memory has been retrieved, sensory information is extracted in order to predict the current observation. Our sensory sensory data is represented in a one-hot encoding, e.g. each element in the vector corresponds to a different sensory experience, and so we model it with a categorical distribution *Cat*. The function *f_x_*(⋯) is 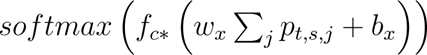, which looks intimidating but in words is simple: we take the retrieved memory, sum over the entorhinal preference cells *j* to produce a vector of the correct dimension. *f_c*_* is a MLP for ‘decompressing’ into the correct input dimensions. The summing over entorhinal preference *j*’ is shown schematically in Figure S4B right, with subsequent decompression transformation, *f_c*_* (⋯), to a one-hot sensory prediction.

#### 2.8 Inference architecture

We have just defined the generative model, however to do anything interesting we need to be able to infer the posterior over the hidden variables. Unfortunately, due to the inclusion of memories, as well as other non-linearities, the posterior *p* (**g***_t_,* **p***_t_ |* **x***_≤t_*, **a***_≤t_*) is intractable. We therefore turn to approximate inference, and in particular the variational autoencoder framework (*15, 45*). Here the inference distribution is parametrised by a neural network, which during training learns how to infer.

The split between inference and generative networks is analogous to the idea of the sleep-wake algorithm. The inference network is ‘awake’ and observes the world, seeing each state as it transitions through the environment. The generative network is used during ‘sleep’ for *training* and where it compares ‘sleep’ generated variables to the inferred ‘awake’ ones. This allows training of *both* networks such that the inference network and generative network learn to align themselves i.e. the generative network learns to predict both sensory data and the variables inferred by the learned inference network (a.k.a recognition distribution) which, in turn, learns to appropriately map sensory events to latent variables.

In defining our approximate recognition distributions, q*_ϕ_*(⋯), we make critical decisions that respect our proposal of map-structure information separated from sensory information as well as respecting certain biological considerations. We use a recognition distribution that factorises as

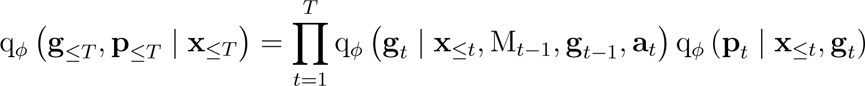

See Figure S2 for inference model schematic. *ϕ* denote parameters of the inference network. The variational posterior can be expressed by the following equations.

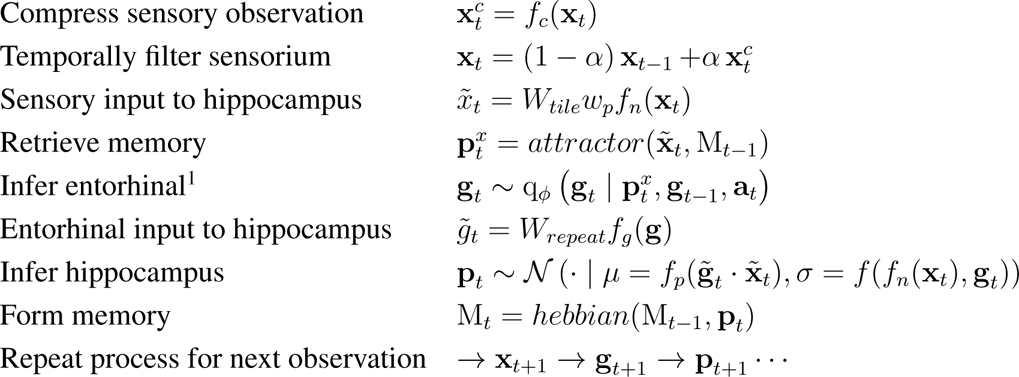

We pictorially show this process (with no Hebbian memory storage) in Figure S5 (just consider the blue stream initially, the second red stream will make sense in Section 2.10). We now explain step by step in words, offering further details and hopefully some intuition.

We take our input **x***_t_*, which is a one-hot encoding of sensory experience (e.g. each element in the vector corresponds to a different sensory experience), and compress it via *f_c_*(**x***_t_*). We compress from a one-hot to a two-hot encoding to reduce the size of the resulting network and ease computation (shown in Figure S4A and Figure S5).

We then smooth this compressed representation over time using exponential filtering. We note that although the exponential smoothing appears over-simplified, it approximates the Laplace transform with real coefficients. Cells of this nature have been discovered in LEC (*55*).

Next, we normalise the representation using *f_n_*() which demeans then applies a rectified linear activation followed by unit normalisation. These representations are then scaled by *w_p_*, and then multiplied by a fixed weight matrix *W_tile_* (which gives the appropriate hippocampal dimension - all dimensions shown in Table S1) to give **x̃***_t_* which is TEM’s sensory input to hippocampus.

We are now ready to infer where we are in the graph. We factorise our posterior on **g** as

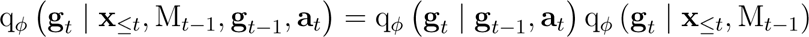

To know where we are, we can path integrate (the first distribution, equivalent to the generative distribution described above) as well as use sensory information that we may have seen previously (second distribution). The second distribution (optional) provides information on location given the sensorium. Since memories link location and sensorium, successfully retrieving a memory given sensory input allows us to refine our location estimate. We use **x̃***_t_* as the input to the attractor network to retrieve the memory associated with the current sensorium, 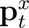. We use MLPs with input 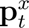 to parametrise the mean and variance of the distribution. This factored distribution is a Gaussian with a precision weighted mean - i.e. we refine our generated location estimate with sensory information.

For Bayesian connoisseurs, we note that, unlike **p***_t_*, these retrieved memories 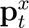 are not random variables in the generative model and are therefore not inferred. Instead they are part of the function in the inference model that learns the approximate posterior posterior on **g***_t_*. Nevertheless they share similarities to **p***_t_*, e.g. they have the same dimension and are pressured to learn similar representations (see Section 3). Biologically, they can be thought of as memories cued only by sensory input, and not inferred from the combination of sensory and structural input.

Now that we have inferred where we are, we are ready to form a new memory - infer our hippocampal representation. After the the entorhinal representation is down-sampled using *f_g_* (⋯), we then multiply by a fixed weight matrix *W_repeat_* (which gives the appropriate hippocampal dimension - all dimensions shown in S1) to give **g̃***_t_*. We define the mean of the inferred hippocampal representation as the element wise multiplication of **x̃***_t_* and **g̃***_t_* followed by an activation function. We choose the *leaky relu* activation function to create sparsity and ensure the only active hippocampal cells are those that receive both map-structure and sensory information. We note that the two fixed weight matrices are designed such that their application, followed by an element wise product between **x̃***_t_* and **g̃***_t_*, is equivalent to an outer product followed by reshaping to a vector (Figure S4).

#### 2.9 Memories

##### 2.9.1 Storage using Hebbian learning

Memories of hippocampal cell representations are stored in Hebbian weights between hippocampal cells. We choose Hebbian learning, not only for its biological plausibility, but to also allow rapid learning when entering a new environment. We use the following learning rule to update the memory:

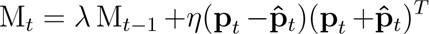

where **p̂***_t_* represents place cells generated from inferred grid cells. *λ* and *η* are the rate of forgetting and remembering respectively. We note than many other types of Hebbian rules also work.

Notably, unlike the generative network, there is no requirement for a memory in the inference network. However, including such a memory allows the network to refine the path integration with landmark information *before* creating its place code and therefore speeds learning dramatically. However, representations in the main paper are observed both in networks that include an inference memory and those that do not.

In networks that do use an inference memory, we can either use the same memory matrix as the generative case (as the brain presumably does), or we can use a separate memory matrix. Best results (and those presented) were when two separate matrices were used. We used the following learning rule for the inference based matrix: 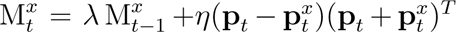, where 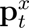 is the retrieved memory with the sensorium as input to the attractor.

##### 2.9.2 Retrieval using an attractor network

To retrieve memories, similarly to (*56*), we use an attractor network of the form

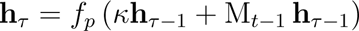

where *τ* is the iteration of the attractor network and *κ* is a decay term. The input to the attractor, **h**_0_, is from the grid cells or sensorium (**g̃***_t_* or **x̃***_t_*) depending on whether memories are being retrieved for generative or inference purposes respectively. The output of the attractor is the retrieved memory.

#### 2.10 Details about embedded hierarchy

Though not a requirement, we embed TEM with the notion of hierarchical scales. TEM abstract location and grounded location (memory) representations, **g***_t_* and **p***_t_* respectively, now come in different frequencies (hierarchies) indexed by superscript 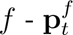 and 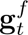. This allows higher frequency statistics to be reused across lower frequency statistics, improving the speed of learning and reducing the number of weights that need to be learnt.

Implementation wise, this means our inference network has several parallel streams of procedure described above, each indexed by the superscript *f*. Each stream has its own learnable parameters (e.g. temporal filtering coefficients in the approximate Laplace transform *α^f^* - a smaller *α^f^* means a longer temporal smoothing window). Otherwise the inference procedure is the same. We schematically show an example of two separate streams in Figure S5. The place where the separate streams can interact is in the attractor network: when storing Hebbian memories, connections from high to low frequencies are set to zero, so that memories are retrieved hierarchically (low-resolution first). These connections are not set to zero when the memory M is used in inference.

The generative procedure is similar to as described above, except for state transitions, where connections in *D_a_* (the action dependent transitions weight from **g***_t−_*_1_ to **g***_t_*) are from low frequency to the same or higher frequency only (or alternatively separate transition streams). There may be different numbers of TEM entorhinal cells in each frequency - this number is described by *n^f^*. When making predictions of sensory experience, we only use 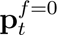 i.e. the highest frequency.

The separation into hierarchical scales helps to provide a unique code for each position, even if the same stimulus appears in several locations of one environment, since the surrounding stimuli, and therefore the lower frequency hippocampal cells, are likely to be different. Since the hippocampal cell representations form memories, one can also utilise the hierarchical scales for memory retrieval.

TEMs hierarchy is consistent with the hierarchical scales observed across both grid cells (*34*) and place cells (*31*), and with the entorhinal cortex receiving sensory information in hierarchical temporal scales (*55*).

### 3 Optimisation

We wish to learn the parameters for both the generative model and inference network, *θ* and *ϕ*, by maximising the ELBO, a lower bound on ln p*_θ_* (**x***_≤T_ |* **a***_≤T_*). Following (*16*) (See (*57*) supplementary material), we obtain a free energy

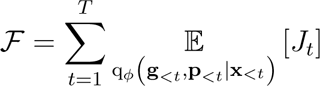

where

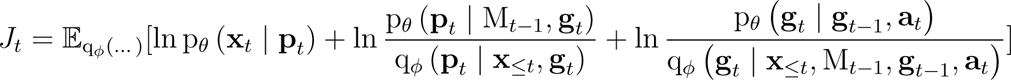

as a *per time-step* free energy. We use the variational autoencoder framework (*15, 45*) to optimise this generative temporal model.

Up to this point the model definition is probabilistic and capable of a Bayesian treatment of uncertainty. However, in the tasks examined in this manuscript there is no uncertainty, so there is no need for this complexity. Hence, we use a network of identical architecture but only using the means of the above distributions - i.e. not sampling from the distributions. We then use the following surrogate loss function to mirror the ELBO:

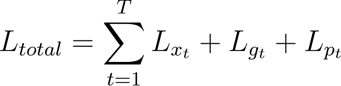

with *L_xt_* being a cross entropy loss, and *L_pt_* and *L_gt_* are squared error losses between ‘inferred’ and ‘generated’ variables - in an equivalent way to the Bayesian energy function.

It is also possible to speed up learning with some augmented losses: As part of *L_xt_* we include losses between sensory experience and 3 different generative model predictions - those generated directly from the inferred **p***_t_*, and also those generated ancestrally through the layers of the network i.e. **g***_t_ →* **p***_t_ →* **x***_t_* and **g***_t−_*_1_ *→* **g***_t_ →* **p***_t_ →* **x***_t_*. When using memory in inference, *L_pt_* includes a memory loss between the retrieved memory and the inferred **p***_t_*.

We use backpropagation through time truncated to 25 steps - this means we roll forwards the inference network for 25 steps, collect the errors and then backpropagate. We then roll forwards the inference network from where we left off etc - i.e. we do not use a sliding window. Longer BPTT lengths are useful as getting an early **p***_t_* wrong will form a bad memory, which then influences the memory retrieval many timesteps into the future. We limit it to 25 for reasons of computational efficiency. We optimise with ADAM (*58*) with a learning rate that is annealed from 1*e −* 3 to 1*e −* 4. Initially we down-weight costs not associated with prediction (*L_gt_* and *L_pt_*). We do not train on vertices that the agent has not seen before. Code will be made available at http://www.github.com/djcrw/generalising-structural-knowledge.

### 4 Simulation and analysis details

All the tasks described below are best ‘solved’ if the underlying structure is learned, even though each structure is different. We now describe the details for the types of graphs we considered, as well as the simulation details.

For all simulations presented above, we use the additional memory module (two separate memory matrices) in grid cell inference. Each time the agent enters a new environment, both memory matrices, M, are reset (all weights zero). Asides from when otherwise stated, the agent randomly diffuses through each environment.

The agent is initially randomly placed in each environment. The agent changes to a completely new environment after a certain number of steps (*∼* 2000-5000 for the 2D graph worlds, lower for smaller environments/ tasks). For 2D graph worlds, typically after 200 *−* 300 environments, the agent has fully learned the structure and how to address memories. This equates to *∼* 50000 gradient updates (1 gradient update per block of 25 steps). For smaller worlds, learning is much faster.

We now describe the dimensions of variables (summarised in S1). We use *n_s_* = 45 (the number of different sensory objects), *n_s*_* = 10 (the compressed sensory dimension) and 5 different frequencies. The number of TEM entorhinal cells in each frequency are [30, 30, 24, 18, 18], and the number of TEM entorhinal cells that project to TEM hippocampus, *n^f^* are [10, 10, 8, 6, 6] (i.e. the first 1/3 entorhinal cells in each frequency). Thus the number of hippocampal cells in each fequence are [100, 100, 80, 60, 60] i.e. *n_s*_* multiplied by each *n^f^*. *λ* and *η* are set to 0.9999 and 0.5 respectively. We note that, like (*27*), a higher ratio of grid to band cells is observed if additional l2 regularisation (penalisation of euclidean norm) of grid cell activity is used.

As mentioned in Section 1, for each task we train on environments of different sizes - this means a true abstract representation must be learned and not just one that is a template map. The learned map must generalise to different sized worlds.

#### 4.1 Transitive inference

The hippocampus is crucial for problems of transitive inference with animals solving novel tasks on first presentation (*11*). And so analogously we test whether TEM can learn about line structures and orderings i.e. if apple is one more than pear and pear is more than monkey, what is 2 bigger than monkey?

To do so we use fully connected graphs, and order the nodes on a line i.e. label each node from 1 to K, where K is the number of nodes in the graph (Figure S1B). Each edge describes an action, e.g. the edge from 5 to 2 describes ‘below by 3’, the edge 4 to 14 describe ‘higher by 10’ etc. This structure and labelling of nodes and edges creates an implicit transitive hierarchy. We use lines of length {4, 5, 6} i.e. number of states {4, 5, 6}).

When TEM navigates these line graph, the actions, **a**, are two dimensional with the first element describing higher/lower and the second element by how much.

#### 4.2 Social hierarchy

The hippocampus is known to be involved in reasoning over social hierarchies (*44*), and again we want to examine whether TEM is capable of learning the abstract set of relationships that govern social hierarchies.

We consider the graph of a family tree (Figure S1C). We limit ourselves to the case where each node has two children. We also eliminate the notion of gender - i.e. aunt/uncle is the same relationship, as is mother/father etc. Each edge corresponds to a family relationship i.e. ‘grandfather of …’. We use 10 types of relationships: {sibling, parent, grandparent, child 1, child 2, aunt/uncle, niece/nephew 1, niece/nephew 2, cousin 1 cousin 2}. We use {3, 4} levels of hierarchy i.e. number of states: {15, 31}.

When TEM navigates these graph, the actions, **a**, are a one-hot encoding of relations such as ‘child of’, ‘grand-parent of’, ‘sibling of’ etc. There are 10 available actions overall.

#### 4.3 2D graphs

The hippocampus and entorhinal system has produced many famous cells, most notably those that have characteristic responses to space (*4, 59*). Thus we consider graphs with spatial properties (e.g. Figure S1D). We consider both 4-connected and 6-connected graphs i.e. those with square or hexagonal symmetries. We use square environments of width 8-11 (number of states: {64, 81, 100, 121}), and hexagonal environments of edge width {5,6,7} (number of states: {61, 91, 127}).

We run simulations in either 6-connected graph worlds, or 4-connected graph worlds. The action is a one-hot encoding - either 4 or 6 dimensional depending on square or hexagonal worlds respectively.

##### 4.3.1 Diffusive behaviour

For diffusive behaviour, the agent has a slight bias for straight paths to facilitate exploration in these larger worlds.

We show all TEM learned entorhinal cells in Figure S6, their autocorrelations in Figure S7, and all hippocampal cells in Figure S8. We note that even in hexagonal worlds TEM sometimes learns hexagonal grid-like cells and sometimes square grid-like cells.

##### 4.3.2 Non-diffusive behaviour

For non-diffusive behaviour (e.g. simulations involving object vector cells), we bias the agents transition behaviours to head towards shiny objects (for object vector cells) or spend more time near boundaries (for border cell representations).

For object-vector cell simulations, we also use an additional distribution in grid cell inference: q*_ϕ_* (**g***_t_ | s_t_*), where *s_t_* is an indicator saying whether the agents is at the location of a ‘shiny’ state. This means that the entorhinal cells can know when it is at a shiny object state. From this, the network builds its own representation encoding vectors from the shiny states. We make one further change to the generative network to encourage the learning of vector representations, by not telling the generative model what action, **a***_t_*, was taken. This encourages it to build representations of what actions will likely be taken (as governed by behaviour). Interestingly, this phenomena is observed in sleep (generative) replay - sequences of head direction activations are divorced from replay of sequences of awake activity locations (*60*).

#### 4.4 Complex spatial tasks

Finally we consider non-spatial tasks embedded in a spatial world. We use the task set-up from (*41*), where rodents perform laps of a circular track. Notably they are only rewarded every 4 laps. Thus the ‘true’ state space of the task is 4 laps not a single lap as space would suggest. This is a non-spatial task (every 4) embedded in a spatial world (circular track). We mimic this task on a loop graph of length *l * n*, with *l* the lap length and *n* the number of laps (e.g. Figure S1E). The sensory observations are identical on each lap, however every *n* laps (i.e. every whole loop of the graph), the first state is a ‘reward’ state - where the reward is a unique sensory observation per environment. We use *n* = 3 laps of length *l* = 12. We increase the backpropagation through time truncation to 60 so that gradient information has access to the whole state space.

We show additional examples of cells that ‘chunk’ as well as those that don’t, from TEM’s hippocampal and entorhinal layers in Figure S9.

### 5 Analysis of remapping data: Preserved place cell-grid cell relationships across environments despite remapping

#### 5.1 Experimental prediction

Our theoretical framework predicts place cells and grid cells retain their relationships across environments - despite place cell remapping - to allow generalisation of structural knowledge encoded by grid cells. More specifically, our framework predicts the following to be true: 1) As has been previously observed experimentally (*28*), our framework predicts that when an animal is moved from one environment to a structurally similar environment but with different sensory experiences, place cells will undergo remapping (e.g. Figure 4C), and grid cells will realign (e.g. Figure 4A). 2) As has also been previously observed experimentally (*28*), we predict the grid cell correlation structure (i.e. relationships between grid cells) within a module will be preserved across environments. 3) Despite realignment and remapping, we predict that, within a grid module, a given place cell will retain its relationship with a given grid cell across environments. For example, if a given place cells firing field is in a given grid cells firing field in one environment, it should remap to a location in a second structurally similar environment that is also in a firing field of that grid cell (Figure S10).

We empirically test for a preserved place cell-grid cell relationship across environments in two datasets from different remapping experiments, in which both grid and place cells were recorded across different environments. We first briefly describe the experimental setup of the experiments, followed by the details of the analyses and results that support our prediction in both datasets. We additionally demonstrate that these results cannot be explained by the place and grid cells not remapping or realigning, and, as has been previously shown (*28*), that the correlation structure of grid cells is preserved across environments.

#### 5.2 Experimental setup

##### 5.2.1 Dataset 1 - Barry et al 2012

In the first dataset (*35*) - dataset 1 - both place and grid cells were recorded from rats in two different environments. The environments were geometrically identical 1*m*^2^ arenas that were in distinct locations in the recording room, and differed in their sensory (texture/visual/olfactory) experiences. Each of seven rats had recordings from both environments in MEC and hippocampal CA1. Each recording day consisted of five twenty-minute trials, in which the rat free foraged in the environments. In between trials the rat was taken out of the arena. Of the five trials on a given day, trials 1 and 5 were in one environment, which the animal is familiar with (having spent at least 100 minutes in the environment), and trials 2-4 were exposures to a second, novel environment. We can therefore test for preserved place cell-grid cell relationships both within and across environments in this dataset.

(*35*) sought to establish the effects of environmental novelty on grid and place cell properties, finding an increase in grid scale and decrease in grid score, as well as an increase in place cell field sizes in novel environments. This effect reduced with exposure to the novel environment over the course of trials 2-4, such that grid and place cells on trial 4 had properties most comparable to those on trials 1 and 5 (*35*). We therefore restrict our analyses of the second environment to trial 4. Further details about the experimental setup can be found in (Barry et al., 2012).

##### 5.2.2 Dataset 2 - Chen et al 2018

We repeat our analyses in a second dataset (*61*) - dataset 2. In dataset 2, both place and grid cells were recorded as mice free foraged in both real and virtual reality environments. These real and virtual environments provide the two different environments for the across environment measures of place cell-grid cell relationships. We do not have a within environment condition for this dataset. As described in full in (*61*), in the virtual reality environment the mice were head-constrained such that head movements were constrained to rotations in the horizontal plane while the mouse runs on a Styrofoam ball. Screens and projectors projected a virtual environment around the mouse and onto the floor from a viewpoint that moves with the rotation of the ball. Hence this system allows expression of free foraging spatial navigation behaviour, analogous to that in the real world.

Both the real and virtual reality environments were square, and size 60*cm*^2^. Trials in the real and virtual environments were 20 and 40 minutes long, respectively. Recordings were made in MEC and hippocampal CA1. (*61*) showed that spatial neuronal cell types that typically characterise 2-dimensional real space, including place cells and grid cells, could be measured in the virtual environment. Of the eleven mice that were trained in the virtual reality system, four had recordings from both place and grid cells, and could therefore be included in our analyses. Further details about the experimental setup and virtual reality system can be found in (*61*).

Details of the number of cells recorded in each animal are found in Table S2.

#### 5.3 Data analyses to test for preserved place cell-grid cell relationship

We tested the prediction that a given place cell maintains its relationship with a given grid cell across environments using two measures. First, whether grid cell activity at the position of the peak place cell activity is correlated across environments (gridAtPlace), and second, whether the minimum distance between the peak place cell activity and a peak of grid cell activity is correlated across environments (minDist; normalised to the corresponding grid scale).

##### 5.3.1 Data pre-processing and critical controls

In the tests presented later, we show results for raw data where we take several steps (with different strictness levels) to avoid possible confounds. Results are shown for all combinations of these choices in Table S3. These include:

###### Defining a grid-score cut-off to ensure entorhinal cells were grid cells

To ensure we are robust to the quality of grid cells entering the analysis, we consider several different grid score cut-offs. We use cut-offs of 0, 0.3 and 0.8. Using less stringent grid cut offs allows more cells and animals into the analysis (Table S2). We would expect our effect to be weaker when reducing the grid score cut off, as the resulting rate maps are likely to be less representative of the grid cell population. Both grid score and scale were computed as in (*35*).

###### Fitting idealised grids to ensure grid-peaks were well-defined

We fit the recorded grid cell rate maps to an idealised grid cell formula (equation 6 from (*62*)), and use this ideal grid rate map to give grid cell firing rates and locations of grid peaks (Figure S11). This leads to a very strenuous control as it ensures that results cannot be driven by *any* differences across grid cells apart from grid phase, grid scale and grid angle (which are the only fitted parameters). This additionally allowed us to use grid peaks that were outside the box. We only fitted idealised grids in situations where we also defined a grid-score cut off (g=0.8) to ensure good model fits.

###### Removing place cells at borders to ensure effects are not driven by border cells

Here we removed all cells whose peaks were *≤* 10% of environment width from the border. The reason we wish to account for border effects is because non-grid MEC cells (such as border cells) rather than grid cells may drive place cell remapping to the borders. We have this criteria for all our analyses.

###### Ensuring cells have actually remapped

Though not data-preprocessing, we ensure that any results could not be confounded by place cells and/or grid cells not remapping/ realigning (i.e. the animal thinking it was still in the same box!). We test this by examining the distributions of spatial correlations obtained when correlating a given place or grid cells rate map in one environment with its rate map in a second visit to that same environment (within environments; only possible in dataset 1) or its rate map in a different environment (across environments). In dataset 1, we found that all the grid cells realigned across environments and the place cells remapped, with spatial correlation coefficients around 0 and distributions similar to those observed in hippocampal global remapping experiments (*28*) (Figure S12). On the other hand, spatial correlations were high upon a second visit to the same environment. Distributions of spatial correlations near 0 for both place and grid cells across environments were also found in dataset 2 (Figure S13). These results suggest that, as expected, grid cells realigned across the environments and the place cells accordingly underwent global remapping; global place cell remapping generally accompanies grid realignment (*28*). That the place and grid cell spatial correlations were near zero means it would be a non-trivial result should the place and grid cell relationship be preserved.

##### 5.3.2 Computing the measures

We first perform the data-preprocessing, making each cell pass the appropriate checks.

We would like to know whether the relationship between place and grid cell pairs is preserved across environments. We propose 2 measures.

###### 1) Does a given grid cell fire similarly at the respective peaks of a given place cell in both environments?

We take a place cell and look at its peak in both environments, which we call P1 and P2. We then take a grid cell, and look at its firing rate at P1 in env1 - we call this X. We look at its firing rate in env2, we call that Y. This gives us a datapoint [X,Y]. We then do this again for the next grid cell, which gives another datapoint. We loop through all the grid cells and place cells for the same animal. Then start again for the next animal. We can then plot all these points on a graph, and find the correlation coefficient - this is the gridAtPlace measure (Figure S14).

###### 2) Does a given grid cell peak at a similar distance from the respective peaks of a given place cell in both environments?

For this MinDist measure, we do the same process as above, but X is now the minimum distance of a grid peak in env1 from P1, and Y is the minimum distance of a grid peak in env2 from P2. We normalise X,Y by grid scale of that grid cell. Note that the minDist measure is only calculated in analyses that fit idealised grids (to cells with grid score 0.8) to ensure that grid peaks are estimated effectively.

For the place cells, we analysed cells defined as place cells in (*35*) and (*61*). Locations of place cell peaks were simply defined as the location of maximum activity in a given cells rate map.

We require each place-grid pair to come from the same animal, but we do not require that the place and grid cells were simultaneously recorded i.e. a place cell may be paired with a grid cell from a different recording session.

Note: If there were only a single grid frequency (or module) in entorhinal cortex, TEM would predict a near perfect correlation across environments between gridAtPlace scores for each grid-cell place-cell pair. However, if either (1) place cells are influenced by phases of more than a single grid module or (2) place cells predominantly received input from a single grid module, but we (the experimenter) do not know which module (as is the case), then we should not predict perfect correlations, only non-zero correlations.

##### 5.3.3 Statistical testing

To test the significance of this correlation and ensure it is not driven by bias in the data, we generated a null distribution by permuting the place cell peak (5000 times) and recomputing the measures and their correlation across trials. We use two possible ways of permuting. Firstly, we choose a position randomly (but still passing our pre-processing steps). Secondly we choose a position from another recorded cell (cells from same and other animals to get enough combinations). We then examine where the correlation coefficient of the non-shuffled data lies relative to the null correlation coefficients to determine its statistical significance. These analyses were carried out separately for both datasets. Again, results from both procedures (for all tests) are reported in S3.

#### 5.4 Which cell types generalise their structure across environments?

As a brief interlude before the main result, we first test whether the correlation structure of each cell type generalises across environments.

##### 5.4.1 Grid cells realign and keep their correlation structure

Indeed, although grid cells realign across environments, their correlation structure is preserved (Fyhn et al., 2007). Although this has been previously demonstrated, we also showed it to be true by demonstrating that the correlation structure between the grid cells was itself correlated (i.e. preserved) across environments. More specifically, we calculated the grid cell by grid cell spatial correlation matrix in one environment, and correlated its upper triangle with that of the correlation matrix in the other environment (a correlation matrix of the same grid cells, but computed in a different environment). We tested this in the single animal with the most recorded grid cells across both environments in each dataset (in a rat with 15 grid cells in dataset 1 [comparing trials 1 and 4], and a mouse with 13 grid cells in dataset 2). This was significant relative to a null distribution generated by permuting grid cell-grid cell pair correlations in both dataset 1 (r=0.55, p¡0.001; Figure S15) and dataset 2 (r=0.95, p¡0.001; Figure S16). These results are expected if the grid cells encode knowledge that generalises across environments. A similar result has previously been reported in (*29*) which included in one of our datasets.

##### 5.4.2 Place cells remap, only weakly retaining correlation structure across environments

We also found this effect to be weakly significant in place cells in dataset 1 (r=0.31, p=0.035; Figure S15) and not significant in dataset 2 (r=0.16, p=0.21; Figure S16).

#### 5.5 Preserved relationship between grid and place cells across environments

Back to our main results in examining whether grid-cell place cell relationships are preserved across environments using our two measures (gridAtPlace and MinDist).

##### 5.5.1 Dataset 1 - Barry et al 2012

As a sanity check, we first confirmed these measures were significantly correlated within environments, i.e. correlated across two visits to the same environment (trials 1 and 5), when the cell populations have not remapped. We see that for both measures there is a significant correlation across the trials (the true correlation coefficient is above 95%of the null distribution of correlation coefficients; Figure S17), for 445 place cell-grid cell pairs. This indicates that upon returning to the same environment, place cells and grid cells have retained their relationship with each other, as expected.

We then tested across environments, i.e. visits to two different environments (trials 1 and 4), to assess whether our predicted non-random remapping relationship between grid and place cells exists. Here we also find significant correlations for all combinations of measures, pre-processing decisions and statistical tests (Table S3). Data for the most stringent/conservative set of inclusion criteria (grid score *>* 0.8, leaving 115 cell pairs) are shown in (Figure S17, gridAtPlace *p <* 0.005, minDist*p <* 0.05)

##### 5.5.2 Dataset 2 - Chen et al 2018

In this dataset, we only have measures for across environments, i.e. visits to the real and virtual worlds. We again found that the gridAtPlace measure was significant across all combinations of measures, preprocessing decisions and statistical tests (Table S3). Here the minDist measure is trending significance (but note this dataset has far fewer cell pairs (Table S3). Figure S18 shows data for the 64 pairs that survived the most stringent inclusion criteria (gridAtPlace *p <* 0.05, minDist *p* = 0.0524)

##### 5.5.3 Remarks

Together, these are the first analyses demonstrating non-random place cell remapping based on neural activity, and provide evidence for a key prediction of our model: that place cells, despite their remapping and grid realignment across environments, retain their relationship with grid cells.

---

**Table S1:** Table showing the number of neurons for each variable. Note the hippocampal dimensions, per frequency, are the multiplication of entorhinal and sensory inputs - coming from the outer product of the two representations

| **Variable** | **Freq 1** | **Freq 2** | **Freq 3** | **Freq 4** | **Freq 5** | **Total** |
| --- | --- | --- | --- | --- | --- | --- |
| Entorhinal $\mathbf{g}$ | 30 | 30 | 24 | 18 | 18 | 120 |
| Entorhinal subsampled $f_g(\mathbf{g})$ | 10 | 10 | 8 | 6 | 6 | 120 |
| Hippocampus $\mathbf{p}$ | 100 | 100 | 80 | 60 | 60 | 400 |
| Temporally filtered sensorium $\mathbf{x}^f$ | 10 | 10 | 10 | 10 | 10 | 50 |
| Compressed sensory observation $\mathbf{x}^c$ | 10 | - | - | - | - | - |
| Sensory observation $\mathbf{x}$ | 45 | - | - | - | - | - |

**Figure S1:**
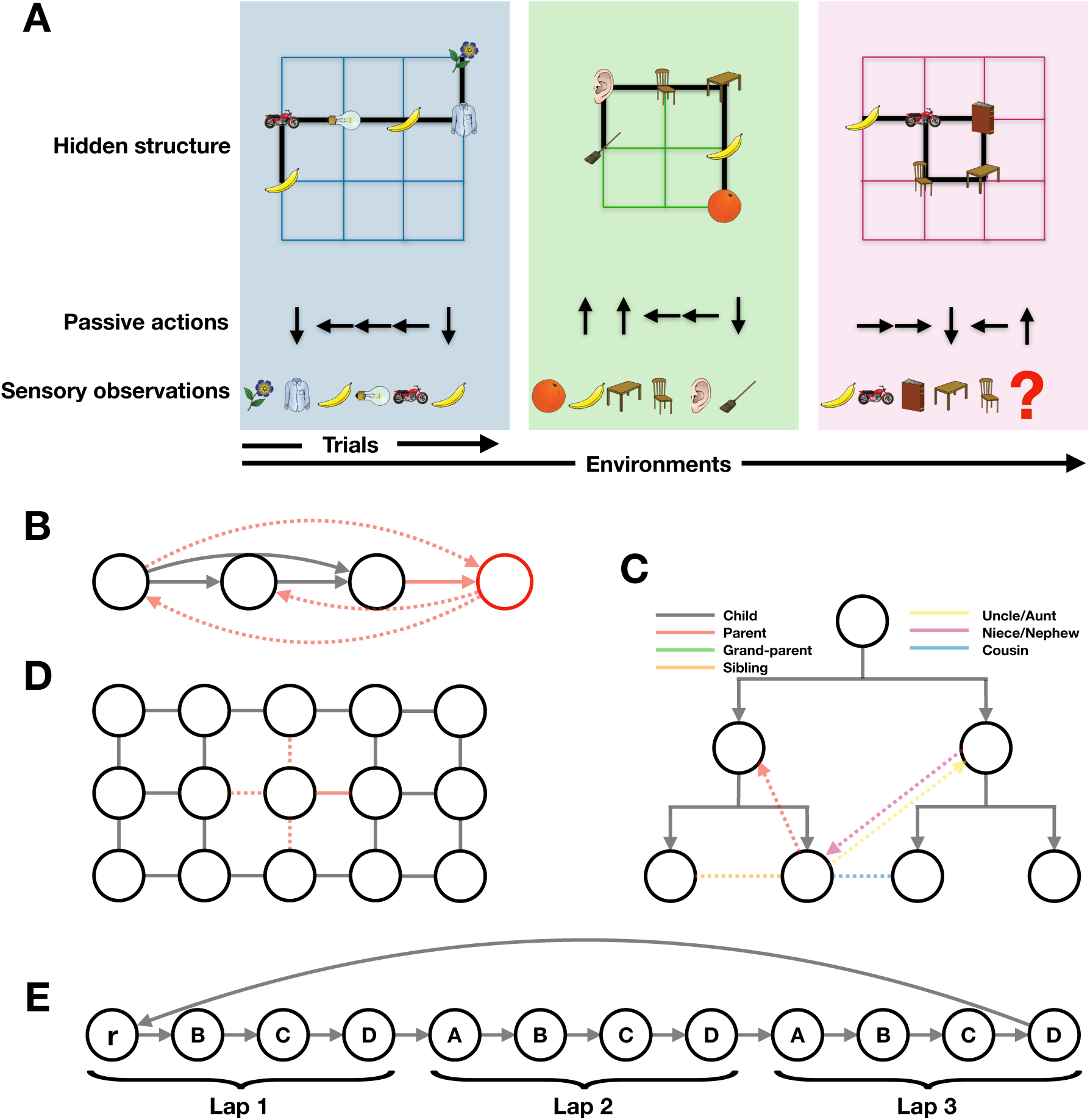
Task schematics for TEM. A) Learning to predict the next sensory observation in environments that share the same structure but differ in their sensory observations. TEM only sees the sensory observations and associated action taken, it is not told about the underlying structure - this must be learned. B) Transitive inference graph. When a new node (red) is seen to be one higher, all other (dotted) relations can be inferred i.e. 3 higher. C) Example graph for a social hierarchy. D) Example graph for 2D structure. E) A complex task embedded in a spatial world. This is a representation of the state space for the task in (*41*). Each lap is of length 4 as the sensory objects (A, B, C, D) repeat every 4 nodes. There are 3 laps in total, and that defines the true state-space as a reward, r, is given every 3 laps.

**Figure S2:**
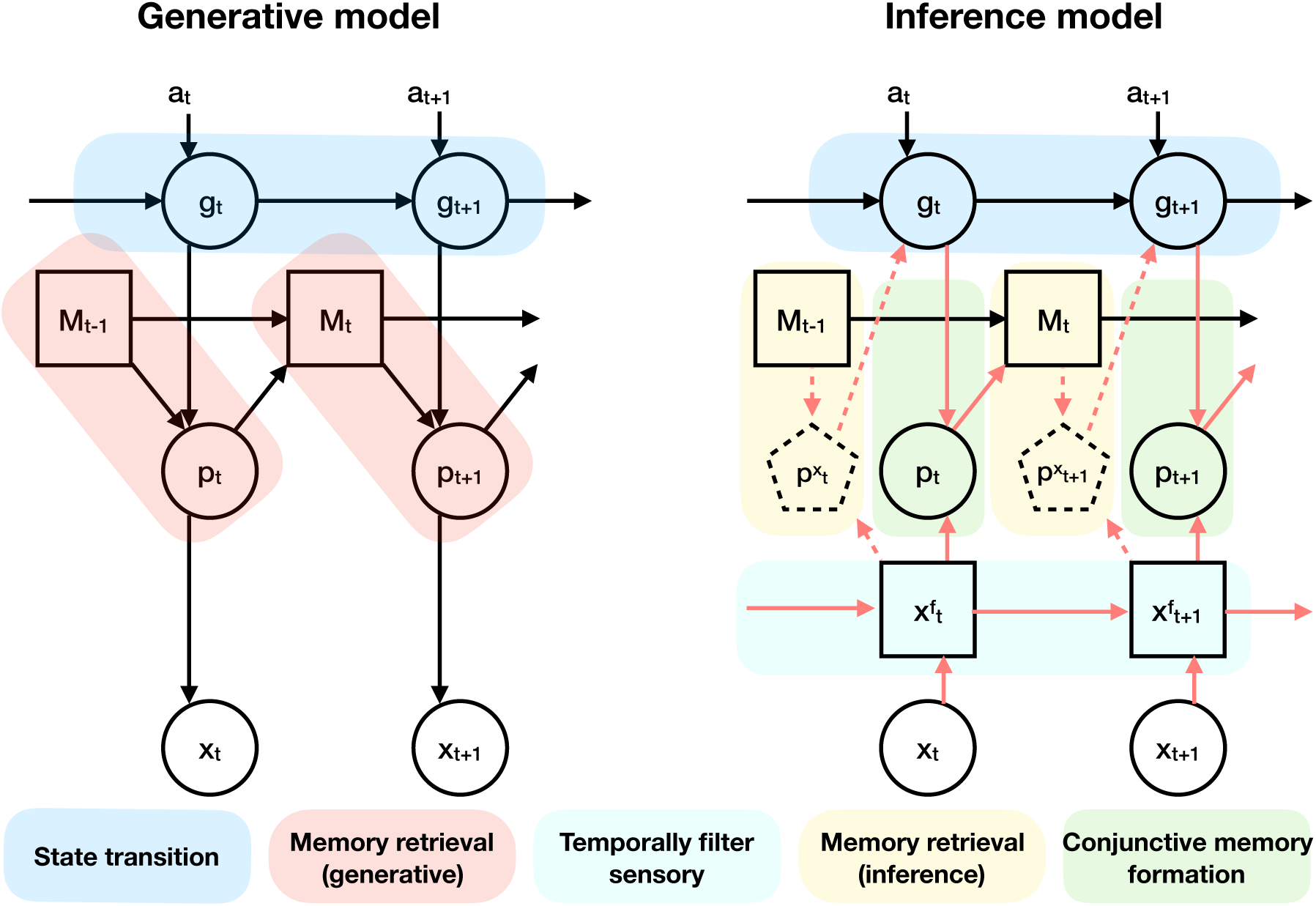
Circled/ boxed variables are stochastic/ deterministic. Red arrows indicate additional inference dependencies. Dashed arrows/boxes are optional as explained in the text.

**Figure S3:**
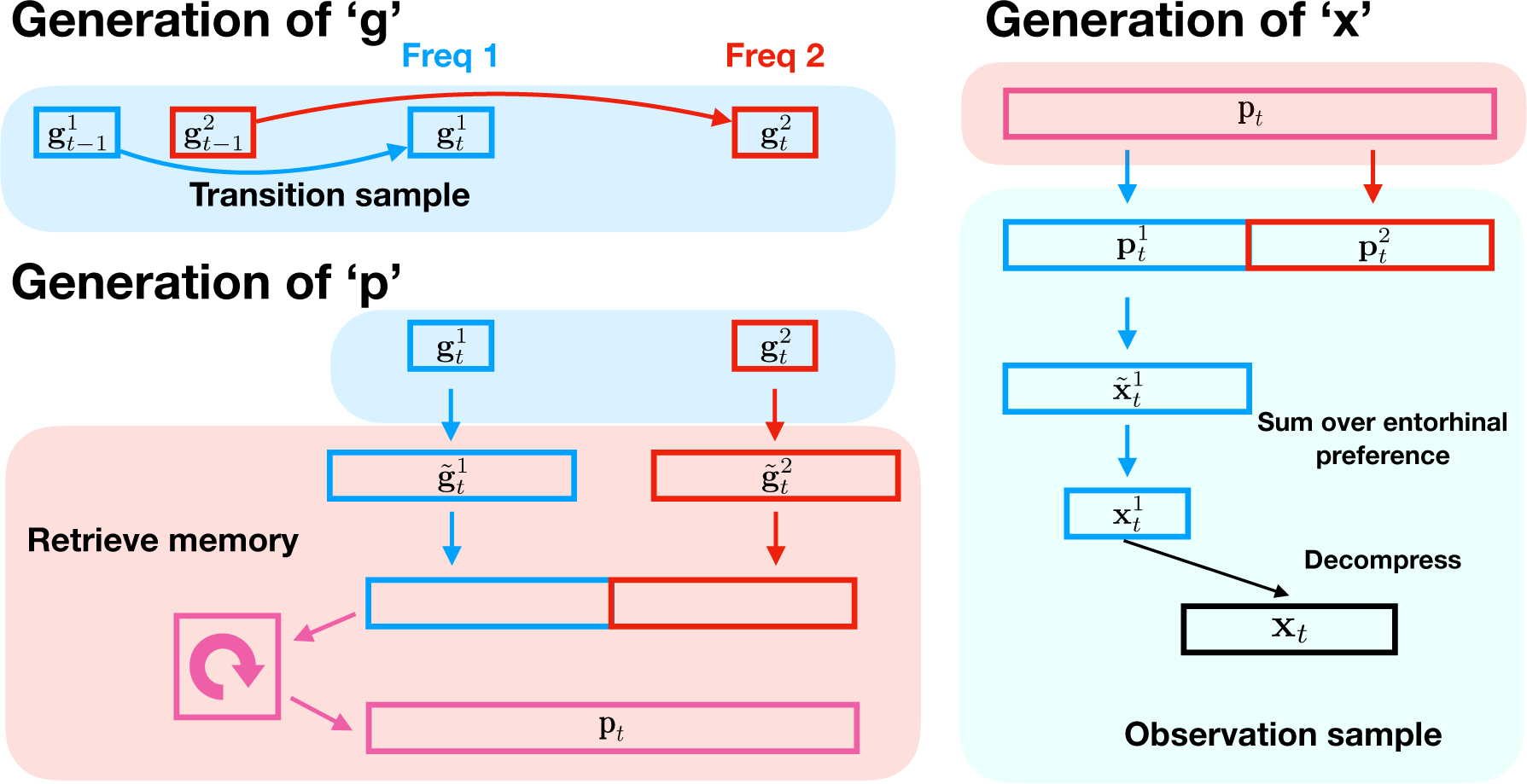
Schematic to show the generative process, making explicit the dimension changes e.g. the sensory decompression step and ensuring the entorhinal input to hippocampus has the appropriate dimension. Red/Blue describe two different ‘frequencies’.

**Figure S4:**
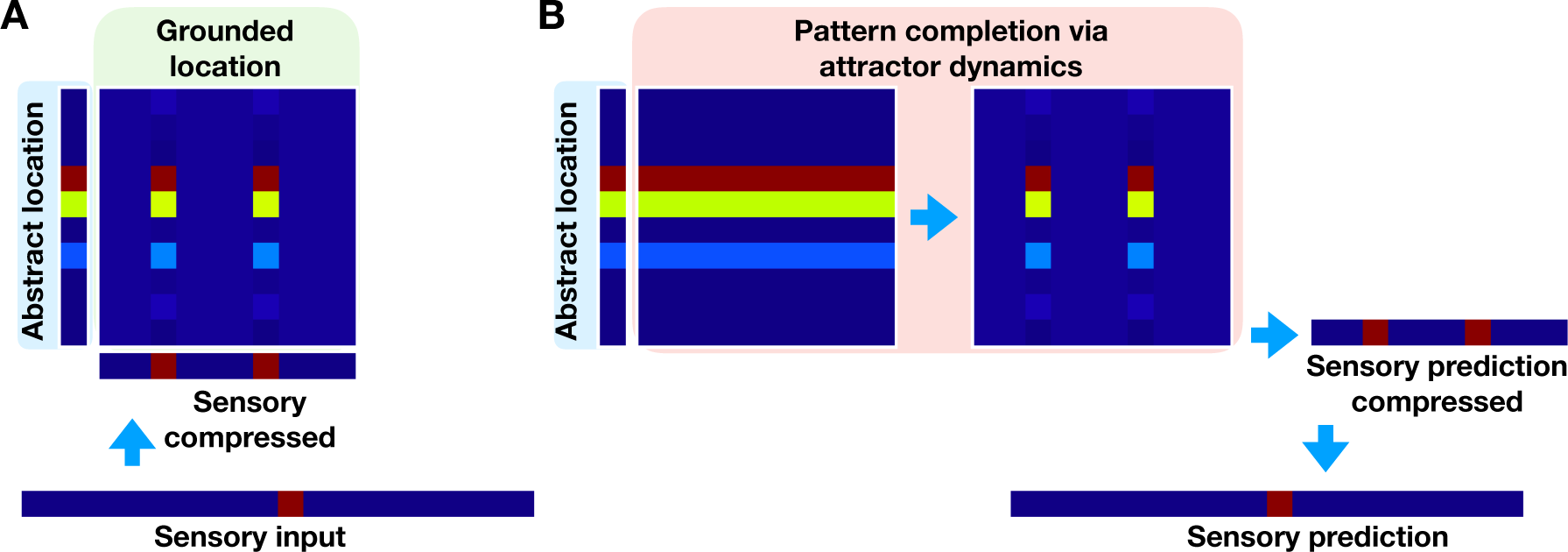
Schematic of conjunction and pattern completion. A) Schematic to show how grounded location (hippocampal) cell representations are formed as a conjunction between abstract location (medial entorhinal) and sensory experience (lateral entorhinal). Note the initial one-hot sensory representation is compressed to a two-hot representation. B) From just abstract location input alone, the correct grounded location representation can be retrieved via attractor dynamics - exciting co-active cells and inhibiting those that were not co-active. The retrieved grounded location can then predict the sensory experience (after summing over the entorhinal preference dimension and then being decompressed). This procedure can also be done ‘the other way round’ with sensory experience alone, in which case the abstract location is predicted.

**Figure S5:**
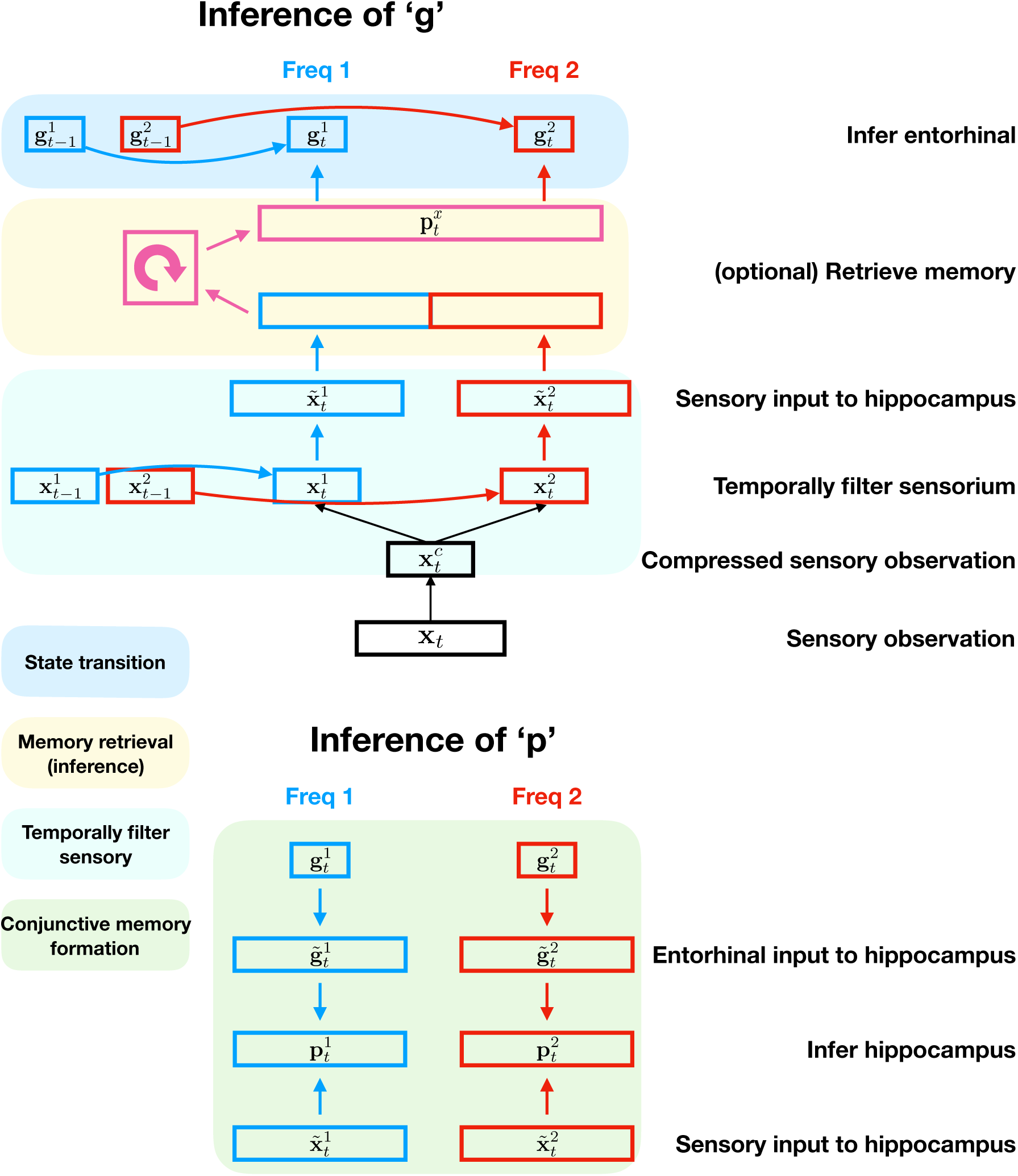
Schematic to show the inference process, making explicit the dimension changes e.g. ensuring the entorhinal and sensory input to hippocampus have the same dimension. We note that this could just be done with a simple linear transformation, but for interpretability we chose to repeat/tile the representations. Red/Blue describe two different ‘frequencies’.

**Figure S6:**
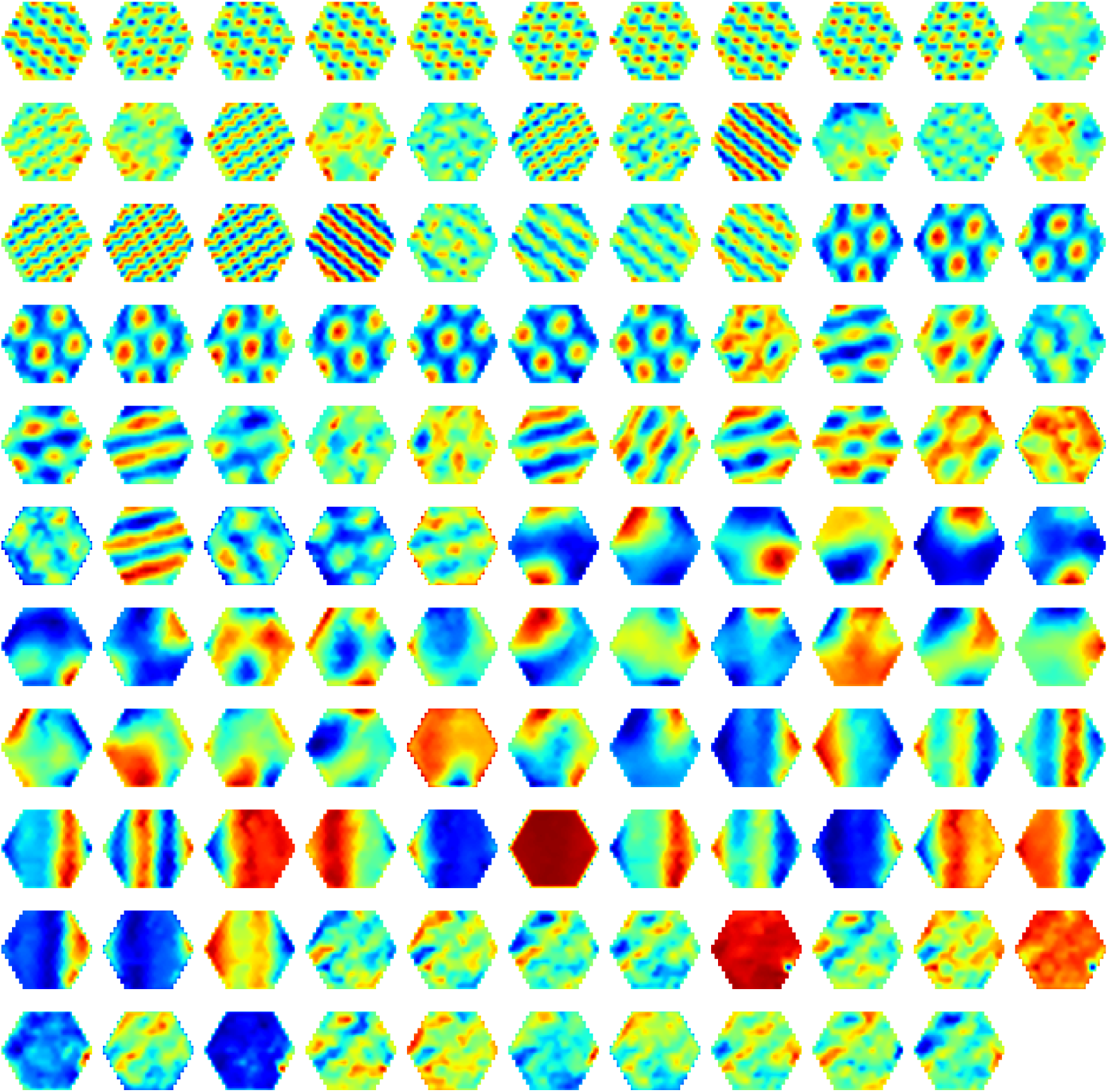
Structural cells, **g**, learned by TEM in diffusive behaviour.

**Figure S7:**
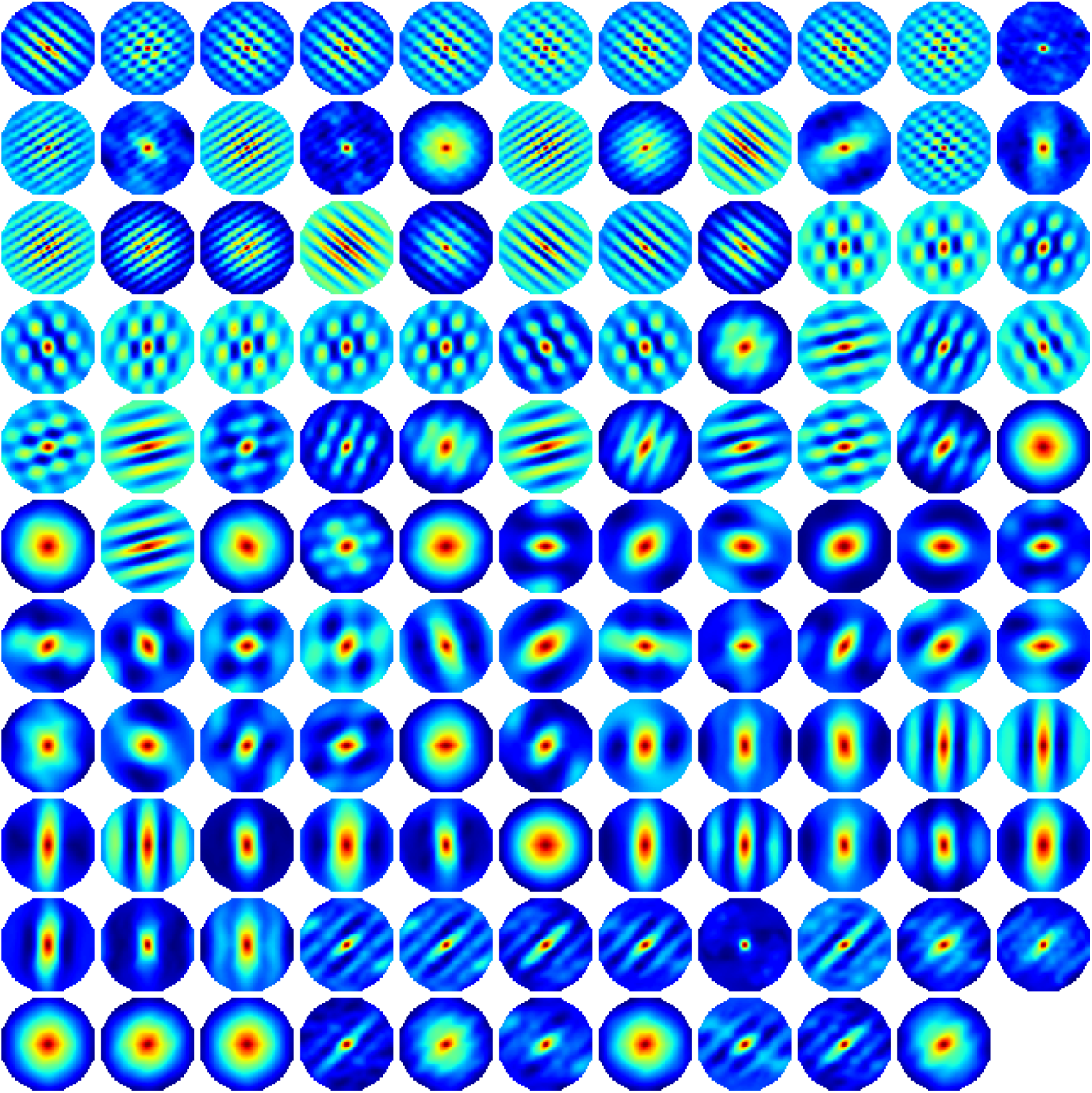
Auto-correlation of structural cells learned by TEM in diffusive behaviour

**Figure S8:**
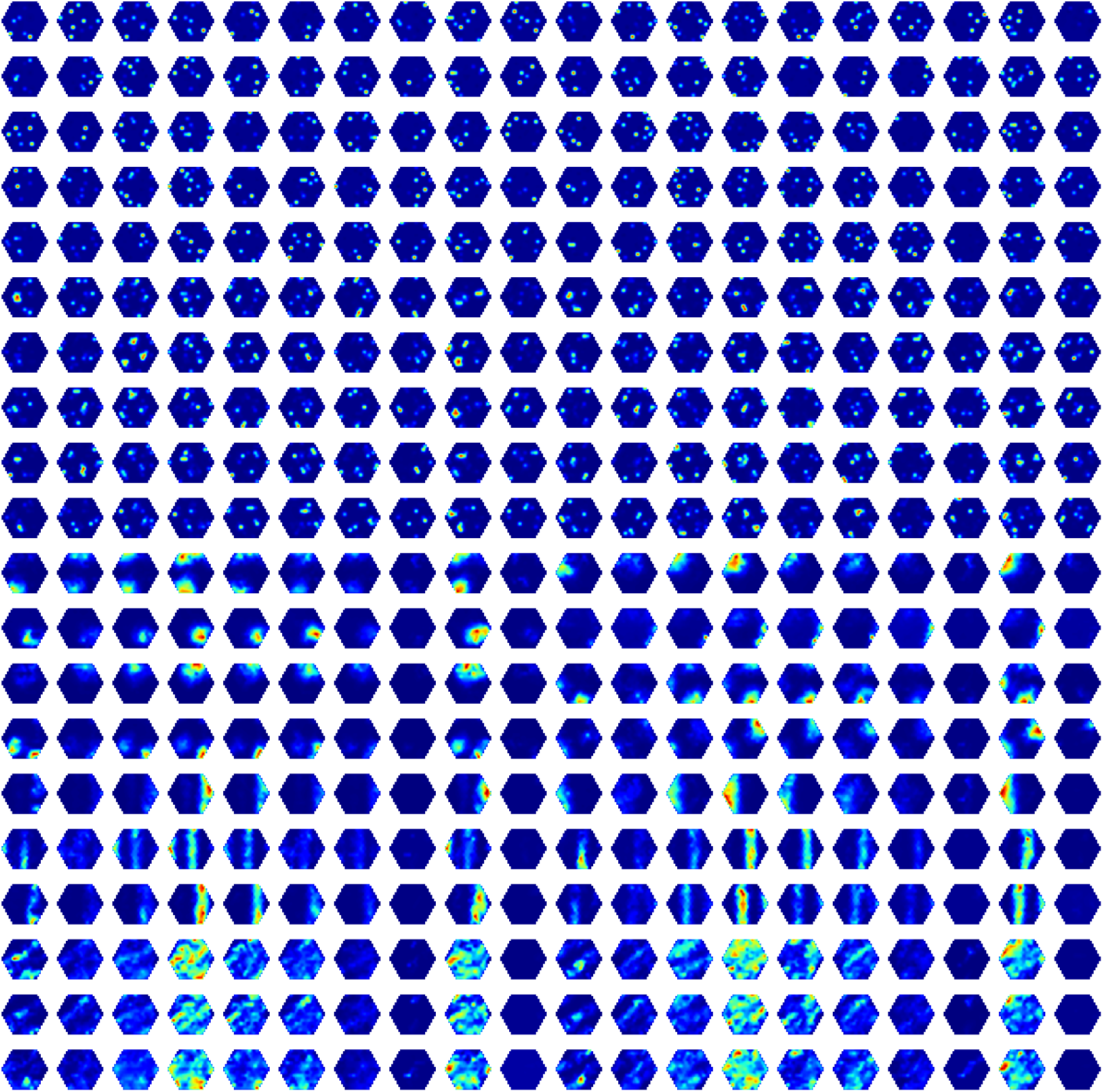
Hippocampal cells, **p**, learned by TEM during diffusive behaviour.

**Figure S9:**
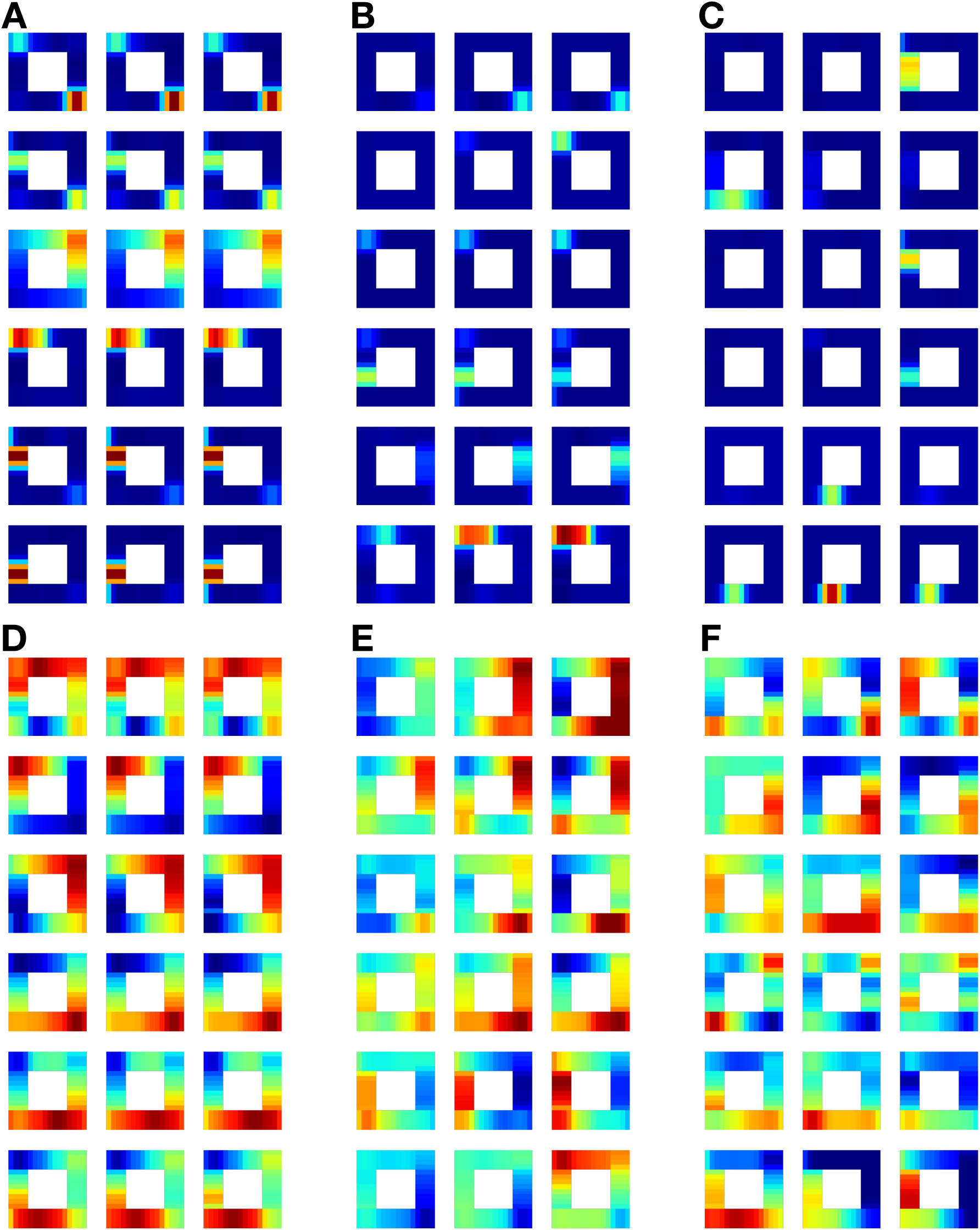
Additional cells rate maps from TEM when performed in the task from (*41*). A-C) TEM hippocampal cells. A) Cells that respond to location on lap - i.e. place cells. B) Cells that appear to ‘count’ the laps. C) Chunking cells. D-F) TEM entorhinal cells. D) Cells that respond to location on lap. E) Cells that appear to ‘count’ laps. F) Cells that chunk the laps.

**Figure S10:**
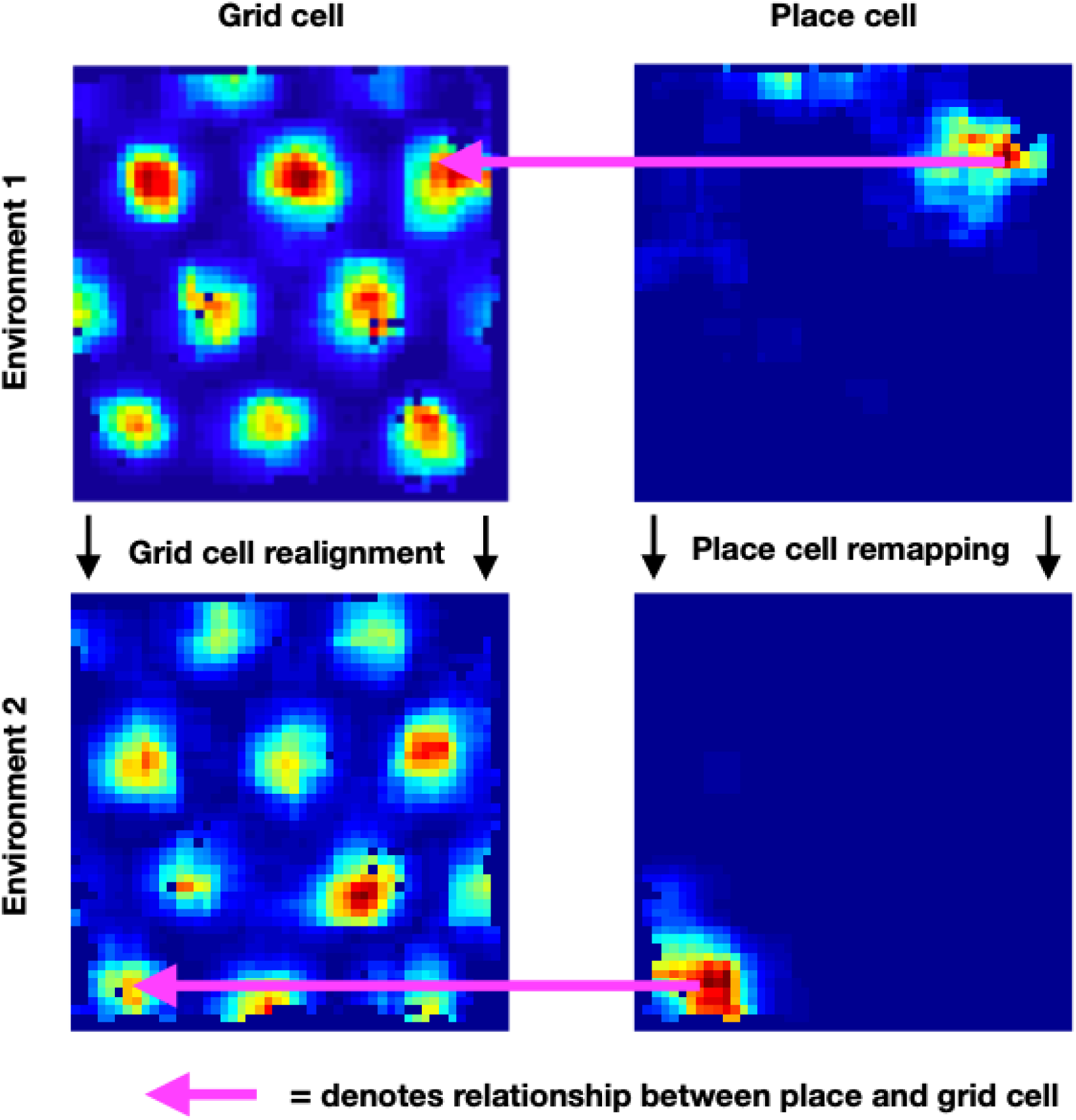
Example place cell-grid cell pair illustrating a preserved relationship across environments. This is an illustrative figure to demonstrate the effect of interest. The rate maps of a place cell and a grid cell in two different but structurally similar environments are shown. The magenta arrow points to the location of the place cell peak in the grid cell rate map for a given environment. It can be seen that in both of the two environments (top and bottom) the place cell peak occurs at a location that is near to the centre of a grid field, despite grid cell realignment and place cell remapping.

**Table S2:** Numbers of animals and cells that enter the across environments analyses. Table showing the total animal and cell numbers that survive the criteria and make it into the analyses where the grid score cut off is ≥ 0.8, ≥ 0.3 and ≥ 0 for both datasets. Note: although the animal numbers may be the same across the datasets, they are different animals.

|  | Animal number | Grid cells | Place cells | Place-grid pairs |
| --- | --- | --- | --- | --- |
| **Grid score $\geq 0.8$** |  |  |  |  |
| **Dataset 1** | 1 | 9 | 12 | 108 |
|  | 6 | 1 | 5 | 5 |
|  | 7 | 1 | 2 | 2 |
| **Dataset 2** | 1 | 5 | 8 | 40 |
|  | 2 | 1 | 6 | 6 |
|  | 4 | 3 | 6 | 18 |
| **Grid score $\geq 0.3$** |  |  |  |  |
| **Dataset 1** | 1 | 15 | 12 | 180 |
|  | 2 | 3 | 10 | 30 |
|  | 3 | 2 | 6 | 12 |
|  | 6 | 3 | 5 | 15 |
|  | 7 | 1 | 2 | 2 |
| **Dataset 2** | 1 | 10 | 8 | 80 |
|  | 2 | 1 | 6 | 6 |
|  | 3 | 2 | 4 | 8 |
|  | 4 | 5 | 6 | 24 |
| **Grid score $\geq 0$** |  |  |  |  |
| **Dataset 1** | 1 | 15 | 12 | 180 |
|  | 2 | 4 | 10 | 40 |
|  | 3 | 2 | 6 | 12 |
|  | 5 | 1 | 4 | 4 |
|  | 6 | 3 | 5 | 15 |
|  | 7 | 2 | 2 | 4 |
| **Dataset 2** | 1 | 13 | 8 | 104 |
|  | 2 | 2 | 6 | 12 |
|  | 3 | 2 | 4 | 8 |
|  | 4 | 5 | 6 | 24 |

**Figure S11:**
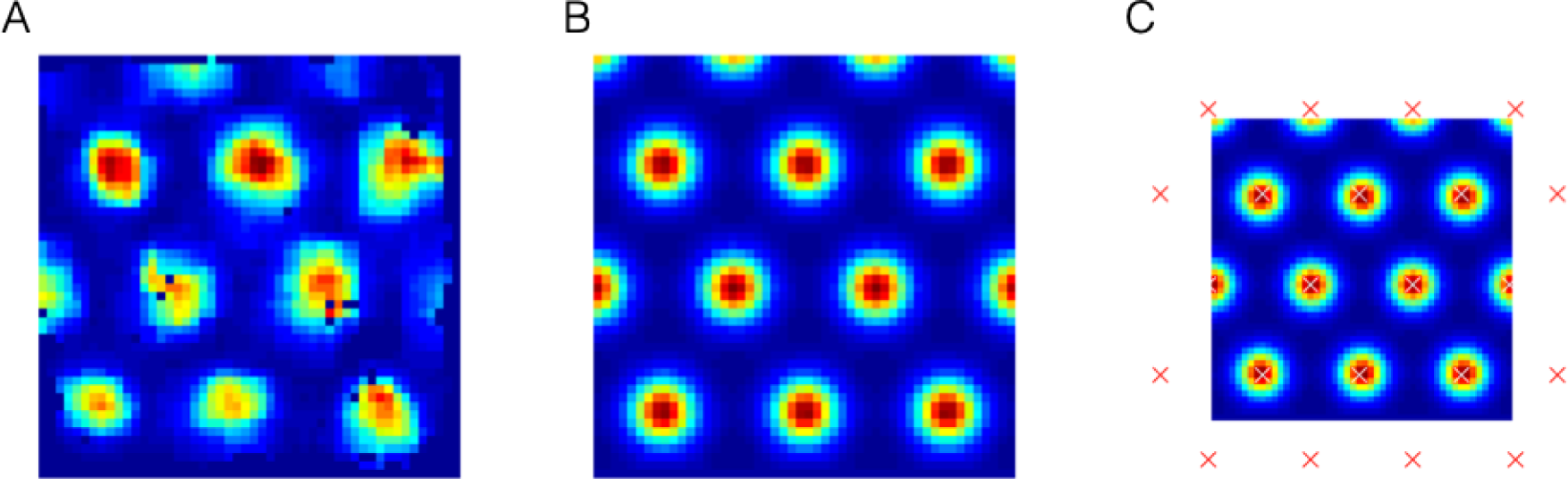
Ideal grid. We fit an idealised grid rate map using the formula from (Stemmler et al., 2015) to the original grid cell rate maps to remove any possible confounds and to ensure that we obtain accurate grid cell peaks. A) An example original grid cell rate map. B) An idealised rate map fit to that in A. C) Accurate finding of grid cell peaks (white crosses) on the idealised grid rate map, which also allows peaks that extend outside the box to be used (red crosses).

**Figure S12:**
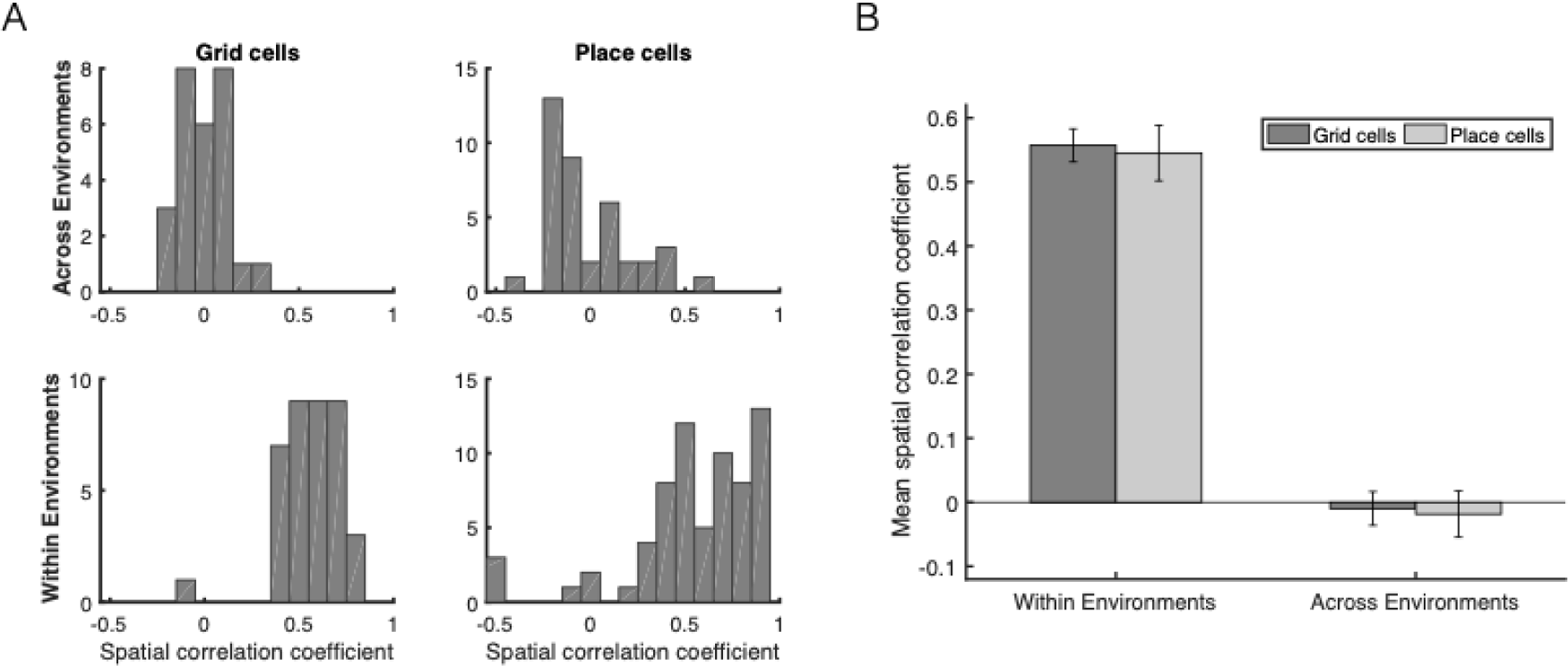
Grid realignment and place cell remapping across environments in dataset 1. A) Histograms showing the distributions of spatial correlations for place and grid cells both within and across environments. B) Bar plots showing the mean (*±* SEM) of these distributions.

**Figure S13:**
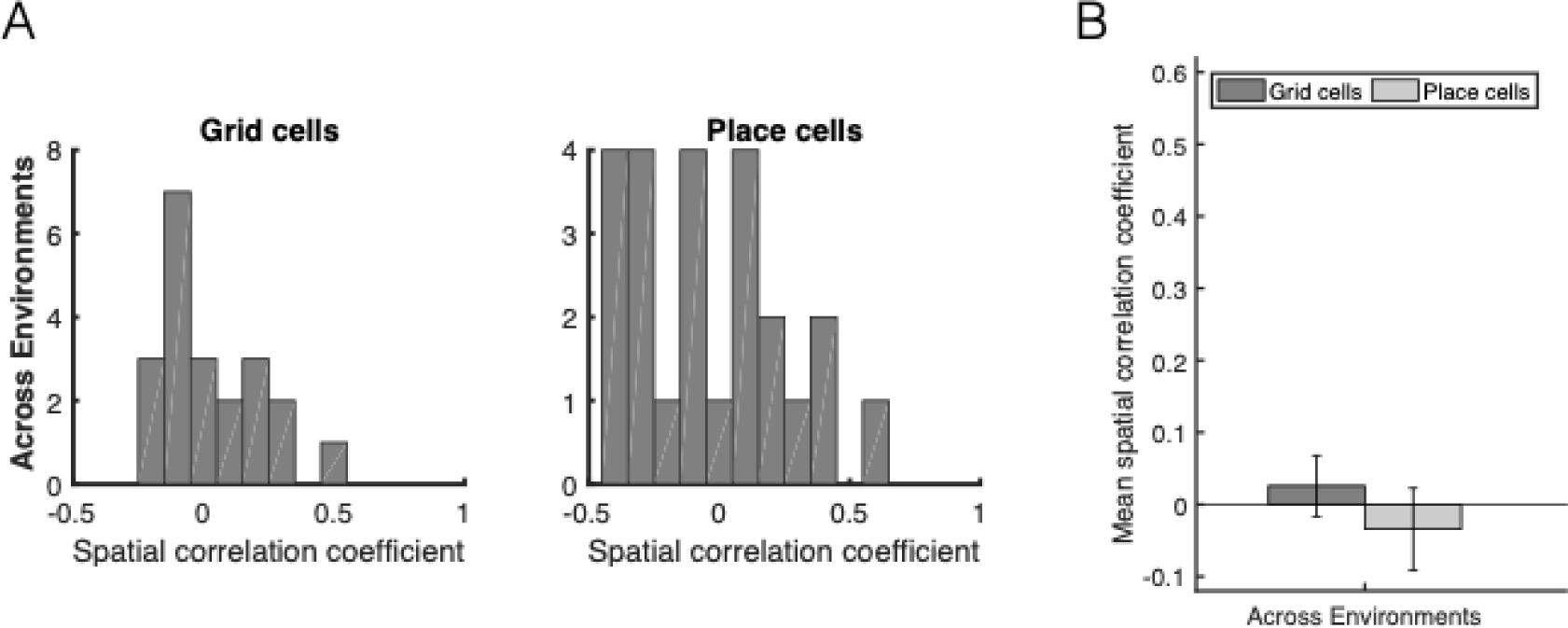
Grid realignment and place cell remapping across environments in dataset 2. The same format as Figure S12, demonstrating distributions of spatial correlations near 0 for dataset 2. Panel B has its axis locked to that of Figure S12B for visualisation.

**Figure S14:**
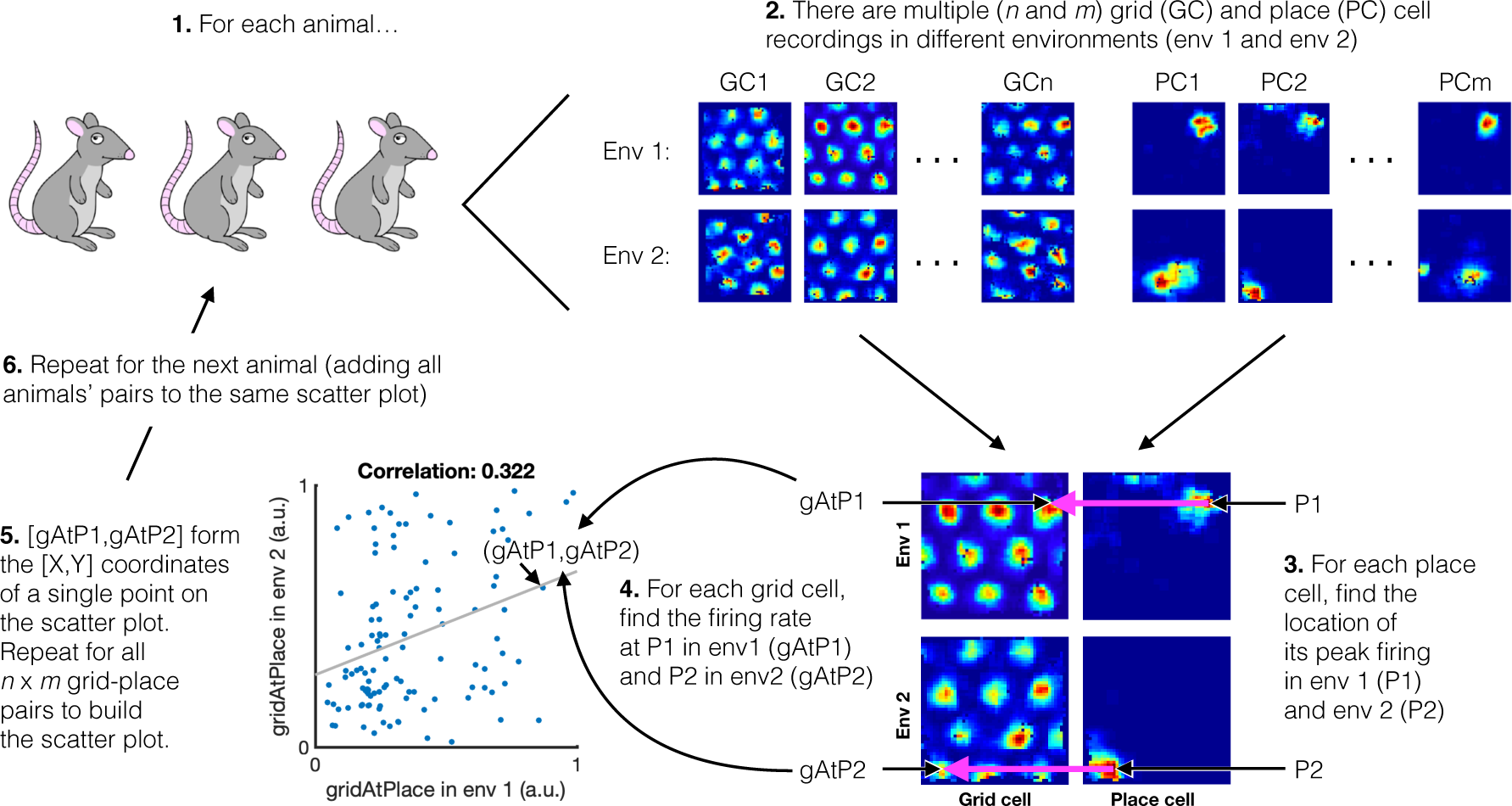
Analysis schematic. Schematic explaining the gridAtPlace analysis. Specifically how the scatter plot is generated. Note that in this figure original grid cell rate maps are shown, rather than ideal grid cell rate maps (Figure S11) that were used to generate the main text figures

**Figure S15:**
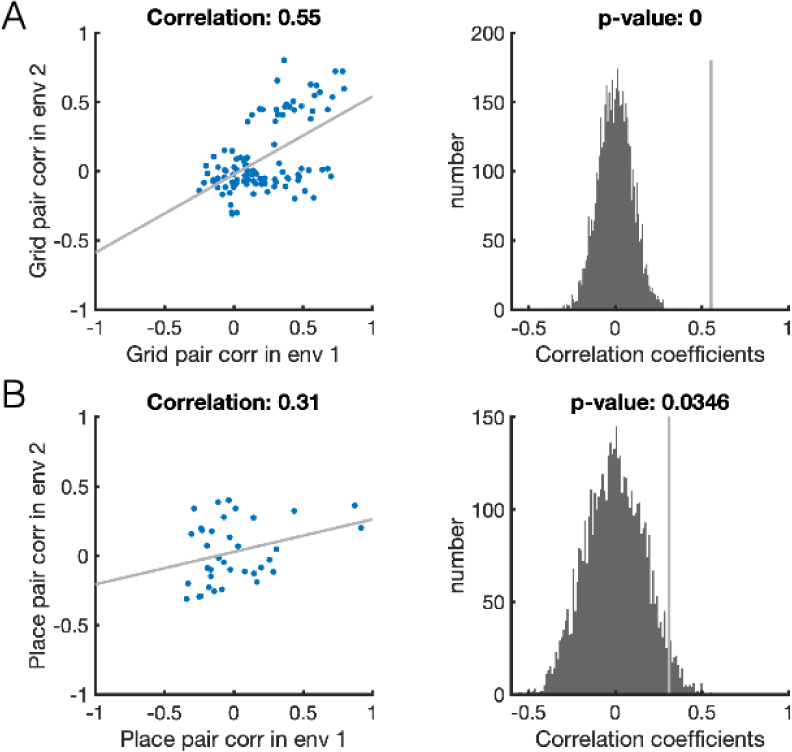
The grid cell correlation structure is preserved across environments in dataset 1. A) Dataset 1. Scatter plot shows the correlation across environments of the spatial correlations of grid cell-grid cell pairs (i.e. the correlation of the upper triangle of two grid cell by grid cell correlation matrices: one from environment 1 and one from environment 2). The histogram shows this correlation coefficient was significant relative to a null distribution of correlation coefficients obtained by permuting grid cell-grid cell pairs. B) Same as A for place cells.

**Figure S16:**
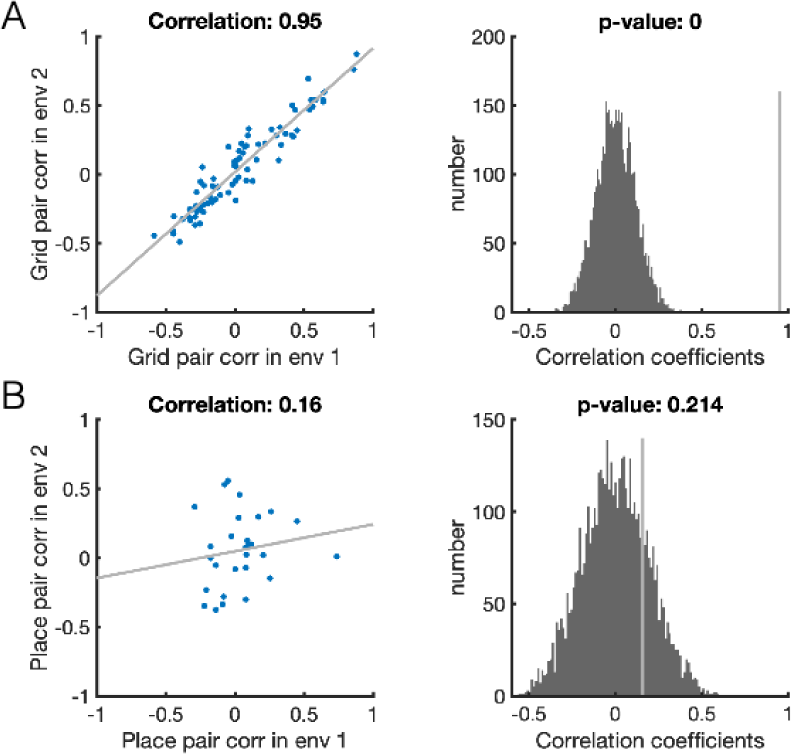
Replication of preserved grid cell correlation structure across environments in dataset 2. A) and B) are the same format as Figure S15.

**Figure S17:**
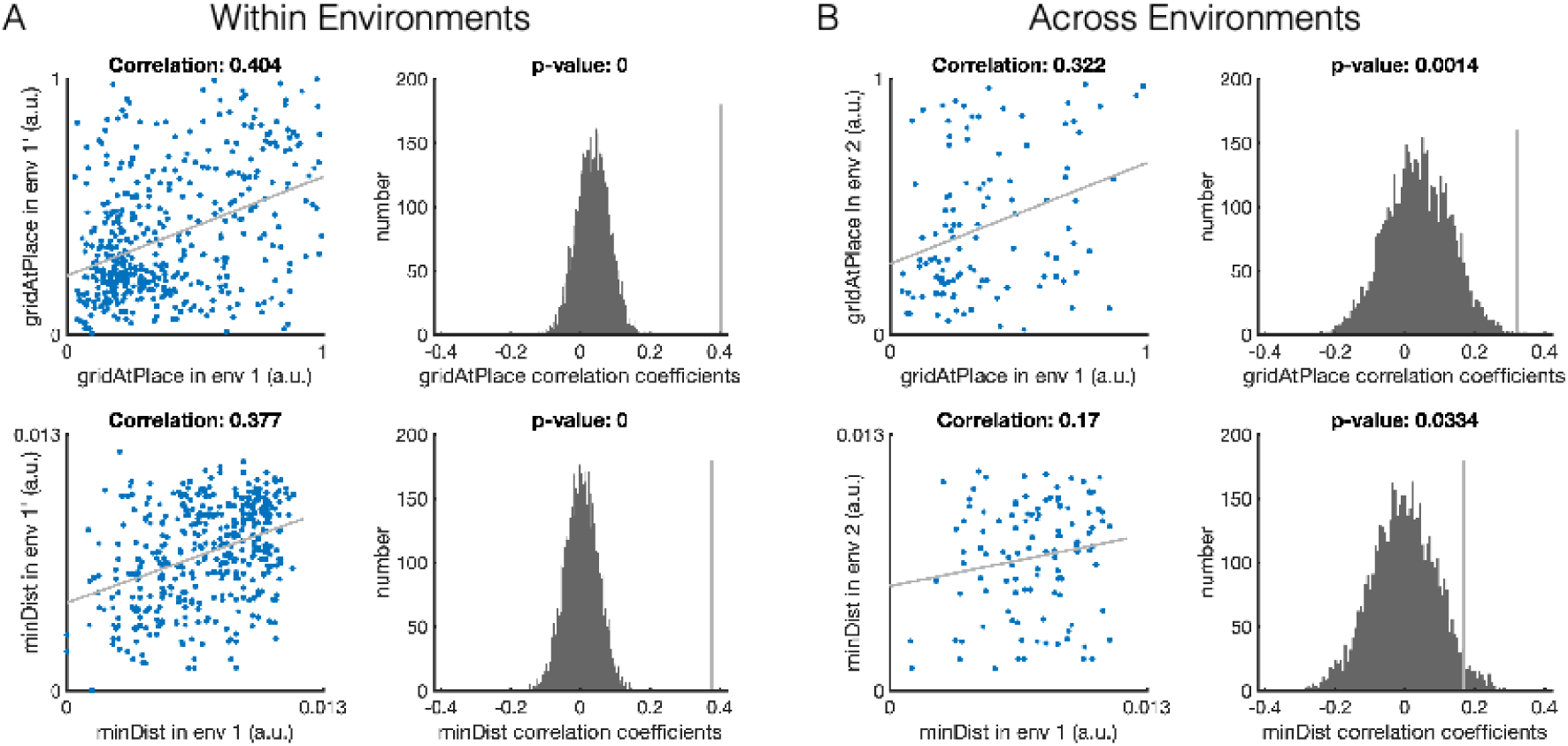
Preserved relationship between place and grid cells across environments in dataset 1. The scatter plots show the correlation of a given measure across trials, where each point is a place cell-grid cell pair. The histogram plots show where this correlation (grey line) lies relative to the null distribution of correlation coefficients. The p value is the proportion of the null distribution that is greater than the unshuffled correlation. A) gridAtPlace (top) and minDist (bottom) measures are strongly significantly correlated over two trials within the same environment, as expected given the same place and grid code should be present. B) These measures are also significantly correlated across the two different environments, providing evidence that place and grid cells retain their relationship across environments.

**Figure S18:**
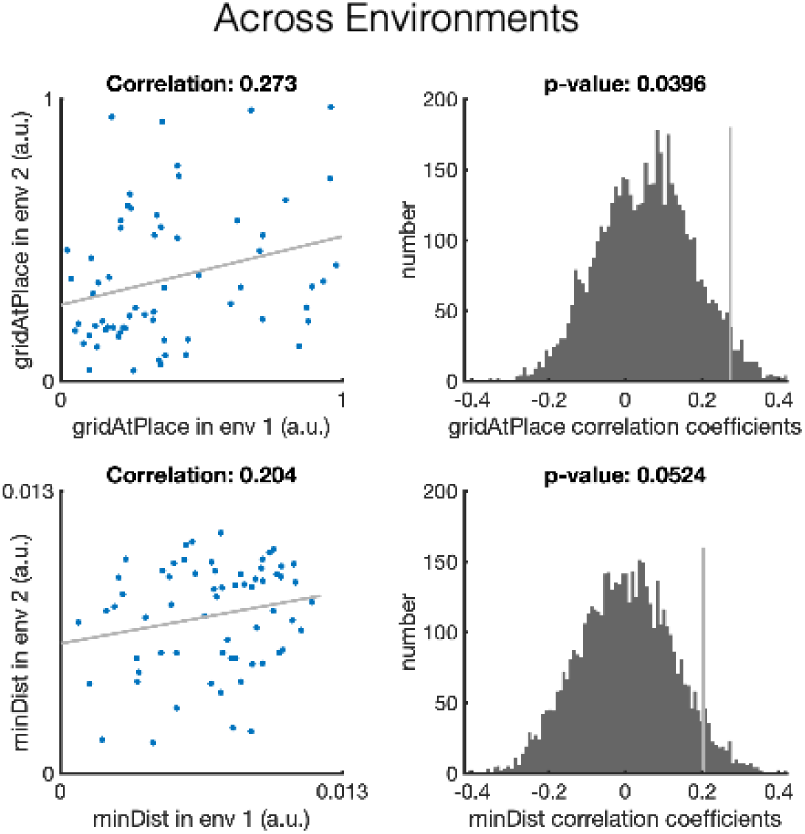
Replication of the preserved relationship between place and grid cells across environments in dataset 2. Figure is of the same format as that of Figure S17. The gridAtPlace measure is significantly correlated at p¡0.05 across real and virtual worlds and the minDist measure is trending very close to significance, replicating the preserved relationship between grid and place cells across environments.

**Table S3:** Analysis results for different parameter settings. Ideal grid 1/0 is whether we used a fitted ideal grid ratemap or not respectively. Permutation type 0/1 is whether we sampled a place cell peaks randomly, or from another recorded place cells. GAP/MD is gridAtPlace and MinDIst respectively. We show the r value and the significant p for both. Results below a 0.05 significance threshold are shown in bold. Our analysis results are robust to parameter settings, with the non-significant results also trending.

| Dataset | Grid cut-off | Ideal grid | Perm type | GAP r | GAP p | MD r | MD p |
| --- | --- | --- | --- | --- | --- | --- | --- |
| 1 | 0.8 | 1 | 0 | 0.322 | **0.0014** | 0.170 | **0.0334** |
| 1 | 0.3 | 0 | 0 | 0.654 | **0.0094** | - |  |
| 1 | 0 | 0 | 0 | 0.630 | **0.0318** | - |  |
| 1 | 0.8 | 1 | 1 | 0.322 | **0.0038** | 0.170 | **0.0352** |
| 1 | 0.3 | 0 | 1 | 0.654 | **0.0356** | - |  |
| 1 | 0 | 0 | 1 | 0.630 | 0.0776 | - |  |
| 2 | 0.8 | 1 | 0 | 0.273 | **0.0396** | 0.204 | 0.0524 |
| 2 | 0.3 | 0 | 0 | 0.554 | **0.0282** | - |  |
| 2 | 0 | 0 | 0 | 0.544 | **0.0404** | - |  |
| 2 | 0.8 | 1 | 1 | 0.273 | **0.0468** | 0.204 | 0.058 |
| 2 | 0.3 | 0 | 1 | 0.554 | **0.0340** | - |  |
| 2 | 0 | 0 | 1 | 0.544 | **0.0468** | - |  |

## Footnotes

1 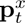 is an intermediary variable retrieved via the memory M*_t−_*_1_ from **x***_t_* - i.e. it represents **x***_≤t_* and M*_t−_*_1_ in the posterior for **g***_t_*.

## Notes

#### Summary of Updates

Removed some typos.

